# Responses to visual motion of neurons in the extrastriate visual cortex of macaque monkeys with experimental amblyopia

**DOI:** 10.1101/2024.07.01.601564

**Authors:** Tom J. Van Grootel, R. T. Raghavan, Jenna G. Kelly, J. Anthony Movshon, Lynne Kiorpes

**Author notes:** Equal contribution (Tom J. Van Grootel, R. T. Raghavan).

## Abstract

Amblyopia is a developmental disorder that results from abnormal visual experience in early life. Amblyopia typically reduces visual performance in one eye. We studied the representation of visual motion information in area MT and nearby extrastriate visual areas in two monkeys made amblyopic by creating an artificial strabismus in early life, and in a single age-matched control monkey. Tested monocularly, cortical responses to moving dot patterns, gratings, and plaids were qualitatively normal in awake, fixating amblyopic monkeys, with primarily subtle differences between the eyes. However, the number of binocularly driven neurons was substantially lower than normal; of the neurons driven predominantly by one eye, the great majority responded only to stimuli presented to the fellow eye. The small population driven by the amblyopic eye showed reduced coherence sensitivity and a preference for faster speeds in much the same way as behavioral deficits. We conclude that, while we do find important differences between neurons driven by the two eyes, amblyopia does not lead to a large scale reorganization of visual receptive fields in the dorsal stream when tested through the amblyopic eye, but rather creates a substantial shift in eye preference toward the fellow eye.

## Introduction

Amblyopia is a developmental disorder in which abnormal visual experience early in life leads to a persistent impairment in many aspects of visual perception. In amblyopia, the eye and retina appear normal; there is no organic disorder at the level of the visual input, it is therefore a disorder of brain development. Amblyopia is commonly characterized as an imbalance between the eyes in visual acuity and contrast sensitivity (Hess and Howell, 1977; Levi and Harwerth, 1977). In severe cases, cortical eye dominance is affected to the extent that there is a complete loss of binocularity and a reduction of the influence of the amblyopic eye in the cortex (see e.g., LeVay et al. (1980); Movshon et al. (1987); Smith et al. (1997); Kiorpes and Movshon (2004b)). Behaviorally, binocular visual processes such as depth perception and fusion are disrupted (see for reviews Birch et al. (2013); Levi et al. (2015)). However, it is also clear that there are deficits beyond these basic visual functions, including diminished contour integration, global form sensitivity, and global motion sensitivity (for reviews see Kiorpes (2006); Levi (2006); Levi (2013); Hamm et al. (2014)).

The neural basis for the sensory losses in amblyopia has been investigated at several stages of the visual pathway (for reviews see Kiorpes (2016); Kiorpes and Movshon (2004b); Levi (2006); Levi (2013)). Movshon et al. (1987), recording at the level of the lateral geniculate nucleus, LGN, found that spatial tuning and neural responsiveness seem unaffected for animals raised with blurred vision. LGN neural sensitivity is similarly unaffected following short and long-term visual deprivation (Blakemore and Vital Durand (1986); Levitt et al., 2001)). The earliest neural correlates of amblyopia are found in V1, where binocularity is disrupted, and where the spatial resolution and contrast sensitivity of neurons driven by the amblyopic eye may be reduced (Kiorpes et al., 1998; Movshon et al., 1987; Shooner et al., 2015; Smith et al., 1997; Wiesel, 1982). In addition, more subtle abnormalities have been reported in neuronal responses in V1 and V2 that may contribute to abnormal binocular interactions and spatial vision (Bi et al., 2011; Hallum et al., 2017; Shooner et al., 2017; Smith et al., 1997; Tao et al., 2014). In general, behavioral losses in amblyopia are more dramatic than the neuronal losses in V1 and V2 based on spatial tuning properties. Thus, it is unclear whether neural abnormalities at the level of V1 and V2 can account for the basic characteristics of amblyopia or the higher-order global deficits documented in psychophysical studies of amblyopia.

Limited data exist on the effect of abnormal visual experience on downstream areas. Studies of strabismic cats found greater disruption of ocular dominance in extrastriate - posteromedial lateral suprasylvian area (PMLS) and area 21a - compared with the striate cortex (Schroeder and Foxe, 2002; Sireteanu and Best, 1992). In the macaque, one serendipitous recording in extrastriate cortex – likely area V4 – showed moderate effects similar to those in V1 (Movshon et al., 1987). Ocular dominance was shifted away from the amblyopic eye, and contrast sensitivity and neural acuity were moderately reduced in amblyopic eye neurons. The only other extrastriate recordings in amblyopic macaques were an extensive investigation of neural tuning properties in area MT, in the dorsal stream (El-Shamayleh et al., 2010).

Although amblyopia is predominantly a disorder of spatial vision, behavioral deficits are also found in motion sensitivity (Hou et al., 2008; Kiorpes, Tang and Movshon, 2006; Knox et al., 2013; Meier et al., 2016; Simmers et al., 2003). MT plays a well-established central role in processing visual motion. It is therefore a likely area in which to identify the neuronal substrate that underlies motion deficits in amblyopes. El-Shamayleh et al. (2010) examined this question in anesthetized, paralyzed amblyopic macaques. In typical MT, neurons are nearly all binocular and respond equally well to stimulation through either eye. In the anesthetized amblyopes, binocularity was strongly disrupted. Analysis of distributions of single unit responses, however, revealed little difference between the eyes in the degree of direction selectivity, preferred speed or motion sensitivity of neurons driven by the fellow and amblyopic eyes. Another property of MT cells, which arises first at the level of MT, is selectivity for pattern motion direction (Adelson and Movshon, 1982; Movshon et al., 1985). Pattern motion sensitivity is a more complex state of visual processing and develops later than typical motion sensitivity in macaques (Hall-Haro and Kiorpes, 2008; Kiorpes and Movshon, 2014), so it might be more susceptible to abnormal visual input during development than direction selectivity. However, again, the distributions of pattern motion sensitivity for the amblyopic and fellow eyes were not different (El-Shamayleh et al., 2010). While these response properties did not differ between the eyes on a cell by cell basis, a population level comparison based on a pooling analysis that accounted for the reduced proportion of amblyopic eye neurons showed that overall motion and speed sensitivity were reduced for the amblyopic eye population. This deficit reflected behaviorally measured losses in the same animals (El-Shamayleh et al., 2010).

In this study, we extended the study of the effect of amblyopia on neuronal response properties in MT by making extensive recordings in *awake* amblyopic macaques. We studied awake, fixating amblyopic macaques to capture the physiological correlates of amblyopia under conditions more like those in which behavioral sensitivity is assessed. We recorded single unit responses to grating, plaid and random dot stimuli in two amblyopic macaques in which substantial contrast and motion sensitivity deficits were documented behaviorally. As in the previous study in anesthetized amblyopes, we found that binocularity in MT was dramatically reduced, and that cells driven by the amblyopic eye were less numerous than those driven by the fellow eye. Responses to amblyopic eye stimulation showed subtle tuning differences, greater response variance and reduced evidence of integration of upstream motion signals when compared to responses driven by the fellow eye. Motion sensitivity, assessed by dot coherence sensitivity, was lower for amblyopic eye cells, independently of the reduced representation of the amblyopic eye. Finally, we replicated some of the key findings from these awake recordings with MT recordings from the same animals under anesthesia. These data extend previous findings from anesthetized macaques and represent the first neurophysiological assessments in the cortex of awake amblyopic macaques. Our results indicate that the changes in visual response properties of cells in amblyopic MT are relatively subtle, suggesting that the main basis for the amblyopic deficit in motion sensitivity is the reduced number and strength of neuronal signals driven by the amblyopic eye.

## Methods

### Subjects

We studied the properties of well isolated single MT neurons recorded from two adult male *Macaca nemestrina*, as well as data from a third age-matched control animal with normal vision. Animal care and experimental procedures were performed in accordance with protocols approved by the New York University Animal Welfare Committee and in accordance with the NIH *Guide for the Care and Use of Laboratory Animals*.

### Artificial strabismus

At the start of the experiment subject GA was 9.5 years, and subject EL 12.2 years old. The left eye of each of these subjects had been surgically deviated at an early age (5 weeks and 3 weeks, respectively), following testing confirming that the animals had normal acuity in each eye. Esotropia (inward deviation) of the eye was created by transection of the lateral rectus muscle; the medial rectus muscle was resected and advanced to the limbus, and the conjunctiva was reattached to the globe. Surgery was carried out under ketamine hydrochloride sedation using sterile surgical techniques. The resulting esotropia was typical in that a reasonable range of motility of the operated eye was maintained. See Kiorpes et al. (1993) for details. Following standard clinical practice, we refer to the treated eye as the amblyopic eye (AE) and the untreated eye as the fellow eye (FE).

### Neurophysiological methods – awake recordings

A stainless-steel head post was attached to the skull as a head-centered reference and to stabilize the head during recordings. In a separate surgical procedure, a stainless-steel recording chamber was implanted over the superior temporal sulcus contralateral to the strabismic (left) eye (as in Zaharia et al. (2019)).

Subjects were trained to fixate a white dot on a grey background, and after a delay received a liquid reward. Trial durations were between 1 and 5 s. We measured eye position with an SR Eyelink 1000 video eye tracker operating at 1 kHz. During the recording experiments the animals had to maintain eye position within a window up to 2 deg diameter for the FE and up to 6 deg diameter for the AE, for which fixation stability was less good.

The subject was seated in a standard primate chair in front of a computer monitor (NEC MultiSync FP2141SB, 22" CRT), with a viewable screen size of 40x30 cm, a resolution of 1280x960 pixels, and a refresh rate of 120 Hz, at a viewing distance of 57 cm at which 1 degree of viewing angle corresponded to 1 cm = 32 pixels. The monitor was calibrated to ensure linear luminance output. For grating and plaid stimuli, the display mean luminance was fixed at a value near 30 cd/m^2^. For stimuli involving dots, the mean background luminance was reduced and the stimuli were 5 deg patches of randomly distributed bright dots, each 0.2 deg in diameter, rendered at a density of 1200 dots/deg^2^/s.

An Apple Mac Pro computer running custom software was used for stimulus presentation, analog input control and experiment sequencing and timing.

Each session began with a brief calibration measurement in which the animal fixated successive, jumping visual targets with the viewing eye to set a linear calibration for eye position.

An epoxy-coated tungsten micro-electrode (FHC, Bowdoin, ME, USA) was introduced daily to record neuronal signals. The impedance of the electrodes was between 1-10 MΩ (measured with 55-70-0 ICM, FHC). These signals were pre-amplified at the head stage, further amplified, and bandpass filtered (300-3000 Hz), using a Dagan 2400A amplifier (Dagan Inc, Minneapolis, MN, USA). The signal was monitored over headphones. Analog neuronal signals were digitized by the computer at 22.05 kHz, with a resolution of 24 bits, and stored on disk with time stamps synced to stimulus timing. The eye signal was recorded via the video eye tracker and resampled by a data acquisition board (National Instruments, NI 6251, 1 kHz) which interfaced with the stimulus computer.

The electrode was sterilized by 20 min immersion in 70% isopropyl alcohol and loaded inside a custom-made stainless steel guide tube, positioned just below the tissue surface (1-2 mm) and held in place by a ABS plastic grid. At the start of the penetration the electrode was lowered using a Kopf 650 hydraulic micropositioner with the FE viewing. We identified area MT from gray matter-white matter transitions and from the isolated neurons’ brisk, direction-selective responses.

After determining the preference of an isolated MT cell with hand mapping, a set of experiments was run with FE viewing, followed by AE viewing. In cases when the isolation was lost during FE viewing, a new MT neuron was searched with AE viewing. During the session the viewing condition only switched back to FE viewing when an AE set was completed fully. This recording strategy biases MT cell encounter rates in favor of the AE.

In the AE viewing condition the FE was occluded. In the FE viewing condition the AE was usually covered. If it was uncovered, both eyes were unobstructed, but the fellow eye was the dominant and fixating eye (FE). Stimulus positions were aligned with the receptive field of the FE. When data were recorded both with and without covering the AE, data obtained with the AE occluded were used in the analyses for the FE condition. No differences were observed when comparing FE responses when the AE was occluded or open.

### Neurophysiological methods – anesthetized recordings

Following the completion of awake experiments in each animal, we made a set of recordings from area MT and the surrounding extrastriate cortex under anesthesia. The methods were similar to those of El-Shamayleh et al. (2010). Briefly, we induced anesthesia with an intramuscular injection of ketamine HCI (10 mg/kg) and diazepam (0.5 mg/kg). We placed catheters in the saphenous veins and intubated endotracheally under isoflurane anesthesia. Anesthesia was then switched to a constant infusion of sufentanil citrate between 6-30 μg/kg/hr for the duration of the experiment. Eye movements were limited by neuromuscular blockade using a constant infusion of 0.1 mg/kg/hr vecuronium bromide. We made a craniotomy and durotomy posterior to the lunate sulcus and angled a micromanipulator 20 degrees anteriorly and ventrally from the horizontal plane. We drove a 32-channel laminar array (Array V32, Neuronexus Inc) to sample activity along the trajectory of the electrode recording in sites in V3, V4t, and area MT. In subjects GA and EL, recordings targeted the hemisphere where awake recordings occurred, while in control subject PF recordings were made from area MT in both left and right hemispheres.

We presented visual stimuli using a Mac Pro running the same software as in the awake-behaving experiments. We presented stimuli separately to the left and right eyes on two CRT monitors (Sony GDM 5402) arranged in a Wheatstone stereoscope configuration. To ensure precise stimulus timing, we used a multi-display adapter (Matrix DualHead2Go) to fuse each CRT display into a single virtual screen at a refresh rate of 85 Hz and a combined resolution of 2048 (1024 + 1024) x 768 pixels. Stimulus parameters were otherwise matched to those used during awake recordings. Signals recorded from the laminar array were amplified and digitized using a 32-channel digital amplifier (Intan Technologies) and written to disk at a sampling rate of 30 KHz via an Open Ephys acquisition system (Siegle et al. 2017).

Following completion of the experiment, the animals were euthanized and transcardially perfused with 4% PFA. Blocks containing recording tracks were sectioned in the sagittal plane in 50-μm increments. Alternating sections were stained for Nissl substance and for myelin using the method of Galyas et al. (1979).

By spike sorting these recordings, we isolated 319 single and multi-units across three recorded animals (GA = 168, PF = 98, EL = 53). Based on the trajectory of the electrodes and response properties at the histologically assigned sites, we localized most recordings in subjects GA and PF to area MT with a small sample of cells recorded in area V3. In subject EL, the brain changes following the extensive preceding chronic recordings rendered isolation of single neurons and unambiguous identification of area MT challenging; therefore, we made a set of recordings in areas V3 and V4t (on the upper bank of the STS).

### Experimental procedures

For the awake recordings, tuning characteristics of MT cells were assessed by running multiple experiments, each with a range of stimulus properties, as summarized in Table 1. Stimuli were fixed at the preferred direction and/or speed once established. Results were analyzed online and in subsequent experiments parameters were optimized as needed. Following hand mapping, the location of the receptive field was determined more precisely by presenting moving dot patches around the identified location. Adjacent circular patches overlapped by 50% on a grid of 3x3 potential locations. To test pattern selectivity, plaid stimuli were constructed by overlaying two identical grating stimuli with 50% contrast separated by 120 deg orientation. The plaid movement direction is at the intermediate angle between the two gratings. Within a given experiment, stimulus conditions were randomly interleaved, with roughly 10% of the trials being blank screen conditions. Stimuli were presented for 250 ms (30 +/- 1 video frames at 120 Hz) per condition, following each other without interruption for the time that the animal maintained fixation. Conditions were repeated in random order for ∼10 repetitions each. For experiments with a large number of conditions (SF and TF combined), at least 3 repetitions were recorded.

**Table 1:** Stimulus conditions.

| Experiment | Stimulus range | Scale | Step size | Number of conditions + blank | Fit |
| --- | --- | --- | --- | --- | --- |
| R.F. loc. | 3x3 patches |  |  | 9 | 2D Gauss. |
| Dot dir. (deg) | 0 - 330 | lin. | 30 deg | 12 + 1 | Von Mises |
| Speed (deg/s) | 0.25 - 100 | log. | $x1.72 \exp(\log(100/0.25)/11)$ | 12 + 1 | Gauss. |
| Coherence (%) | 0 - 100 | lin. | 10% | 11 + 1 | Weibull |
| Grating dir. (deg) | 0 - 330 | lin. | 30 deg | 12 + 1 | Von Mises |
| Plaid dir. (deg) | 0 - 330 | lin. | 30 deg | 24 (plaid & grat.) + 2 | Von Mises (dir.) |
| *SF and TF separate:* |  |  |  | 11 + 10 + ( 1 + 1 ) |  |
| SF (c/deg) | 0.1 - 10 | log. | $x1.58 \exp(\log(10/0.1)/10)$ | 11 + 1 | Gauss. |
| TF (c/s) | 1 - 31.6 | log. | $x1.47 \exp(\log(11.5/1)/9)$ | 10 + 1 | Gauss. |
| *SF and TF combined:* |  |  |  | 6 x 6 = 36 | 2D Gauss. |
| SF (c/deg) | 0.25 - 8 | log. | x2 | 6 |  |
| TF (c/s) | 0.5 - 16 | log. | x2 | 6 |  |

Abbreviations: *R.F.*: Receptive field, *loc.*: location, *dir.*: direction, *Gauss.*: Gaussian distribution

We used a spike-triggered method to locate visual receptive fields: the stimulus patch boundaries at every spike event were added to estimate the extent of the receptive field. Both visual latency and the measured eye position were taken into account. This yielded a gaze-centered probability map which was subsequently fitted with a two-dimensional Gaussian (see below) from which receptive field location and size could be derived. For experiments other than receptive field mapping, firing rates were averaged per condition.

In experiments on anesthetized animals, we assessed the tuning characteristics of neurons differently to accommodate multineuron recordings. Rather than fixing the preferred direction or speed, we ran a full series of drifting dot stimuli at all combinations of 8 directions and 5 speeds. We also ran a full series of drifting gratings at all combinations of 12 orientations and 8 spatial frequencies. The receptive field mapping procedure was the same between awake and anesthetized experiments, except that in anesthetized experiments, we presented three fields of dots on each trial at different locations. The stimuli were typically presented on a 10x10 grid that spanned a larger enough area of the visual field to capture the receptive fields of all the neurons recorded across the laminar array.

### Data analysis

*Latency.* Response latency was calculated using the method of Smith et al. (2005), in which the stimulus and data stream are time shifted to identify the maximum point of cross-correlation, which is then taken as the latency. Most latencies were determined from dot direction experiments. In other cases, latency was obtained from experiments in the following order of priority: 1) grating direction, 2) plaid direction, 3) dot speed and 4) receptive field experiments.

*Eye dominance.* Eye dominance was obtained by regressing firing rates recorded when viewing with the AE or FE. Firing rates for all matched stimulus conditions were compared, i.e., the computation was not restricted to “best” conditions. We fit a line taking account of the variability of the response measurements through the mean firing rates in each eye. The slope of the line determines the ocular dominance index (ODI).

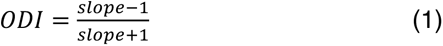

When the firing rate through one eye did not change with that from the other eye, the fit was a straight line with a slope of 0 (horizontal) or ∞ (vertical), yielding an ODI of –1 or +1. AE preferring cells had ODI <= 0, and FE preferring cells had ODI > 0. Figure 5A shows this analysis for 3 example neurons.

For cells that failed to respond through one eye, the ODI was determined from hand mapping experiments. When no spikes were observed, ODI was scored with an integer value of +1 (no spikes from stimulation of the AE) or –1 (no spikes from stimulation of the FE). When the cell’s spontaneous firing was reduced during hand-mapping, indicating suppression, the cell was manually scored with ODI of +1.5 or –1.5. When a cell was responsive during both viewing conditions but responded to different stimuli through the two eyes, the cell was scored with ODI = 0. If a cell was only tested with one viewing condition (e.g. only FE) an ODI could not be obtained and the cell was grouped under the tested viewing condition in population analyses. Note that these are distinct from ‘manually scored’ cells.

#### Distributional comparisons

Unless otherwise stated we performed permutation tests (10,000 permutations) to determine the likelihood that the medians of neuronal population measures differed. Depending on the a priori assumption, the test was either performed one-tailed or two-tailed, which is indicated in the figures and text (one-tailed: ‘>’ or ‘<’, two-tailed ‘<>’).

#### Response variability

We used the method of Goris et al. (2014) to estimate the reliability of neuronal firing. This method fits the relationship between the mean and variance of spike counts with a quadratic that models the modulation of the mean spike count (μ) by multiplying with a fluctuating gain variance 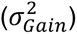:

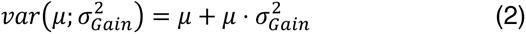

where the mean is calculated from recorded spikes during several trials over the full length of stimulus presentation. This parameter is a measure of reliability and can be compared between different neuronal populations or viewing conditions.

#### Neuronal motion sensitivity

To obtain a coherence threshold from neurons, the electrophysiological coherence-response function was converted to a neurometric function (Britten et al., 1992; El-Shamayleh et al., 2010). As applied in earlier work, this technique creates receiver operating characteristics (ROCs) to measure neuronal *discrimination* performance for opposite directions. Here we used similar methods to measure coherence *detection* performance by comparing neuronal activity at 0% coherence with coherent motions. Such detection thresholds are typically comparable to discrimination thresholds (see El-Shamayleh et al. (2010), Figure 6).

To simulate a psychophysical response based on a population response we pooled the activity of multiple neurons (El-Shamayleh et al. (2010)). Different pool sizes were evaluated (see supplementary figure 1), but for the main results we used a pool size of 10 neurons. The total spike count was collected from randomly selected trials of 10 randomly selected (without replacement) cells for all stimulus conditions. A decision rule compared spike counts in trials with dots moving in the preferred direction with spike counts during zero coherence trials. This was repeated 1000 times for both animals, both viewing conditions, and all coherence levels, resulting in a simulated population psychometric response as function of coherence. Thresholds were obtained by fitting a cumulative Weibull function through neurometric coherence responses. As is conventional, the threshold was taken to be the coherence value corresponding to a value of 0.82 of the fit.

In some cases we extrapolated beyond 100% coherence to estimate threshold. We do not consider coherence thresholds *> 120%* and these extrapolated thresholds are not included in the distribution comparisons.

#### Neuronal tuning

We derived relevant tuning parameters by fitting the descriptive functions described in Wang and Movshon (2016) to neuronal responses.

Spatial and temporal frequency (SF and TF) tuning were determined in two experiments. In one, either SF or TF was kept constant and tuning was determined for the other parameter separately. In the other, SF and TF were both varied, interleaved, in a single experiment. If this latter experiment was run, the cell’s preference was obtained from that data set.

#### Pattern motion selectivity

Pattern motion selectivity can be determined by calculating partial correlation coefficients (Movshon et al., 1985), Fisher transformed to Z-scores (Smith et al., 2005). A pattern index is computed by taking PI = Z*_p_* – Z*_c_*. where Z*_p_* and Z_c_ are the Z-transformed pattern and component correlation coefficients. The resulting PI values are used to classify the cells as pattern selective (PI > +1.28), component cells (PI < –1.28), or unclassified (–1.28 > PI < 1.28) (Smith et al., 2005).

#### Two-dimensional data

We approximated two-dimensional data – including receptive field location and size, spatial and temporal frequency tuning, speed and direction tuning for moving dots, and spatial frequency and orientation tuning – using two-dimensional Gaussian functions. We took the peaks of these approximations as optimal values, and took the full-width at half-maximum as tuning bandwidths, expressed below as receptive field size, 2D velocity tuning, and 2D spatial frequency tuning. These bandwidth measures can be thought of as the square roots of the area of the half-maximum level set of the 2D Gaussian fits.

## Results

### Psychophysics

Both animals underwent behavioral testing prior to the start of the physiological experiments, to characterize their spatial vision and motion perception. Behavioral testing was conducted using standard two alternative forced choice methods (see Kiorpes et al., 2006; Kozma and Kiorpes, 2003). Their visual performance on two tasks measured with each eye is shown in Figure 1. In both animals, spatial contrast sensitivity was attenuated for the AE (in red) across a wide range of spatial frequencies. Motion sensitivity was assessed by measuring coherence thresholds in a motion direction discrimination task using random dot kinematograms. Subject GA showed a dramatic deficit in sensitivity across a broad range of speeds, while Subject EL was impaired only for the slow speeds. The latter pattern is one that we previously reported to be typical of amblyopic macaques (Kiorpes et al., 2006) and is also seen in human amblyopes (Meier et al., 2016). This pattern of deficits can be explained by poor sensitivity to fine spatial scales that is characteristic of amblyopia, which also limits the perception of slow movement.

**Figure 1.**
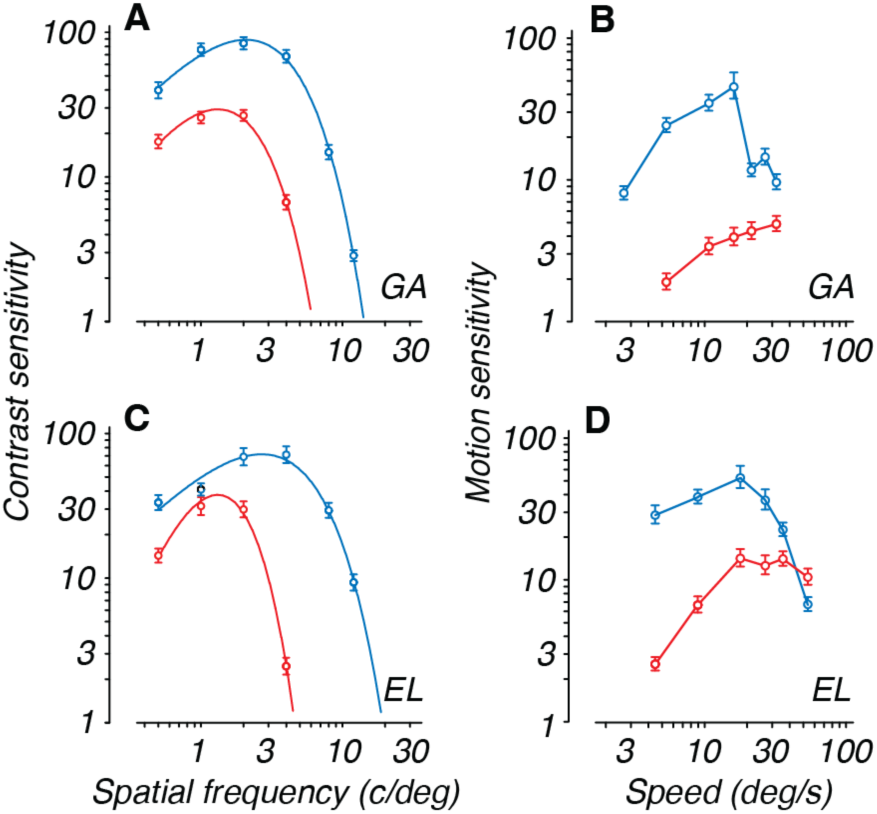
Spatial contrast sensitivity (left) and visual motion sensitivity (right) for the two amblyopic monkeys. Sensitivities measured with FE are shown in blue, AE in red. Bars indicate standard errors. **A, B**: subject GA, **C, D**: subject EL. **A and C** show contrast sensitivity measured with stationary gratings of different spatial frequencies. A double exponential function is fitted through the mean contrast sensitivity data. The amblyopic contrast sensitivity deficit is most prominent at high spatial frequencies. **B and D** show motion discrimination performance measured with moving dots of variable coherence at different speeds. The amblyopic motion sensitivity deficit is most prominent at low speeds.

### Eye position

Throughout each awake behaving recording session, the animal maintained fixation on a dot on the screen. Stimuli were aligned with the cell’s receptive field by changing the stimulus position and/or fixation position on the CRT screen. Figure 2A shows the precision and accuracy of fixation for each eye of both animals. The fixation position during FE viewing was highly accurate on average (left panels, blue). Fixation precision was higher in subject EL than in GA, even with the FE. The red crosses show fixation accuracy for the AE. On average, eye position was biased rightward (nasalward) of the fixation dot. This agrees with the observation that both animals exhibited eye position drift towards the right when viewing monocularly with AE. The animals made temporalward catch-up saccades in the direction of the fixation dot to compensate for the drift displacement, similar to nystagmus. Subject GA exhibited mostly horizontal drift movements, while EL showed moderate drift both horizontally and vertically. Reduced precision is reflected in the larger standard deviations in Figure 2A.

**Figure 2.**
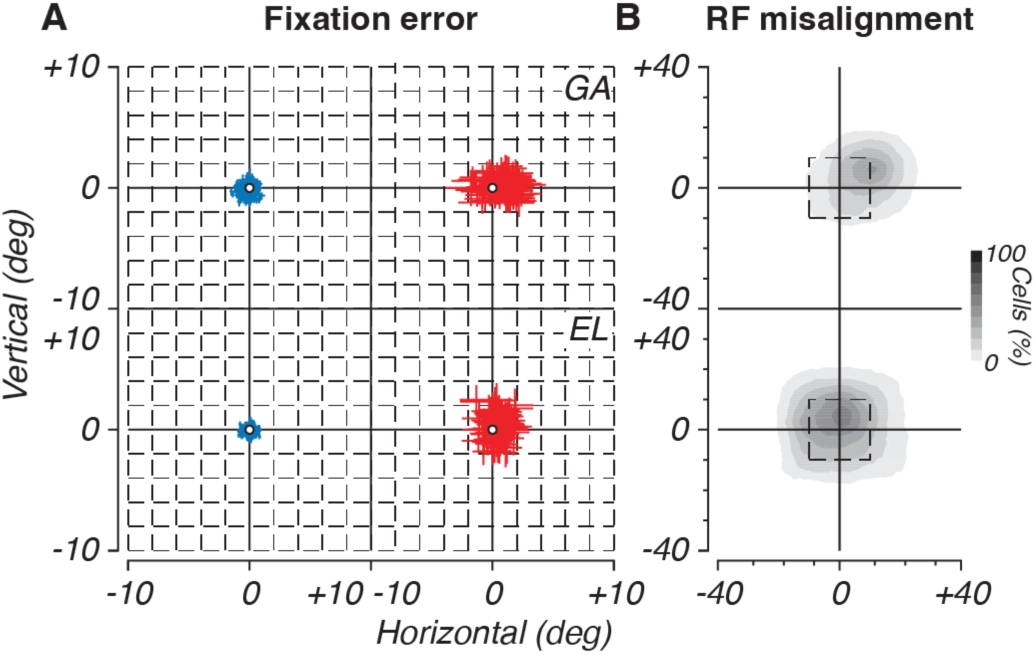
Fixation accuracy and precision, and the effect on measured receptive field location. Top panels subject GA, bottom panels subject EL. **A.** The center of the panels aligns with the location of the fixation dot. Every colored cross indicates the average fixation location throughout the recording of a cell. The length of the crosses are standard deviations of the horizontal and vertical misalignments. These values are obtained by accumulating the deviation from the fixation dot throughout the recording. The resulting distribution was fitted with a 2 dimensional Gaussian. **B.** Apparent misalignment of the obtained receptive field locations: The gray scale reflects the number of cells that have overlapping receptive fields for AE viewing. For every cell the receptive field is obtained during AE viewing by reverse correlating neuronal firing with stimulus position (see Methods). The activity is normalized such that cells receives equal weights. The AE receptive field is aligned with the location obtained during FE viewing (center of panel). The dashed box indicates the scale of panel A. The deviation from the origin indicates that on average the receptive fields are misaligned, by an amount that is larger than expected from the reduced fixation precision indicated by panel A.

The systematic fixation errors posed a problem for aligning the stimulus to the receptive field. As a result, the stimulus location that maximally drove the same cell in the two eyes needed to be shifted by a systematic and substantial amount (Fig. 2B). Interestingly, this shift was much larger than the fixation error: tens of degrees instead of within a degree (note the difference in the axis scales between panels A and B of Fig. 2). The receptive field shift was similar in magnitude to the angle of the strabismus (GA: ∼30 deg, EL: ∼20 deg). This might indicate that there was a substantial remapping of receptive fields. An alternative explanation is that subjects were fixating eccentrically, that is, they did not align the fixation spot with their fovea. An off- fovea fixation location could also give rise to observations shown in Figure 2B.

In the anesthetized recording experiments, the AEs were aligned with the stimulus using a reversible ophthalmoscope to determine the location of the foveas. The receptive fields were not shifted from their expected – aligned – locations, suggesting that as we had suspected, the monkeys did not choose to fixate with their foveas when viewing through the AE.

### Awake physiological recordings

A total of 1436 cells were recorded across the two animals; 389 cells were removed from the analyses as not visually responsive or categorized as not being in MT (some were judged to be in MST, for example, on the basis of receptive field size and location). An additional 30 cells were discarded because isolation was poor. A total of 1017 MT cells were used for further analysis, 606 from GA and 411 from EL. Of those, 410 cells were only recorded under one viewing condition (AE only or FE only; 268 from GA and 142 from EL); 607 cells were recorded under both viewing conditions (338 (GA) and 269 (EL)). 21 cells from GA were run under FE viewing both with and without occluding the AE. The cells in this category allowed us to assess the effect of occluding visual input to the AE during FE viewing. There was no systematic effect on the neural activity, so trials under FE viewing conditions with and without occlusion of the AE were combined. Note that when both eyes were open, the FE was always the fixating eye.

Because isolation was occasionally lost part of the way through the full set of experiments, not all stimulus conditions were run on all cells. We collected data for dot direction tuning in 86% of cells; the receptive field mapping experiment in 71%; a dot speed experiment in 86%; dot motion coherence sensitivity measurements in 53%; grating direction tuning in 48%; and grating spatial and temporal frequency tuning in 46% and 45%, respectively.

### Responsiveness and example tuning

Our tuning analysis depended on the estimated visual latency, which we calculated using the cross-correlation method of Smith et al. (2005). Figure 3A shows that for both eyes and both animals the latency is within the typical range of MT neurons (50-100 ms). Some latency estimates are very long or short (latency < 30 ms or > 150 ms; extreme bins). This is likely the result of low shared variance across conditions, perhaps due to an overall low firing rate, which makes the latency estimate difficult. The medians of each distribution, and statistical tests, are listed in *Table 2: Latency*. When the medians of the populations, excluding the outliers, are compared, AE cell latencies (red) were significantly *shorter* than FE cell latencies (blue). We had expected latencies in AE cells to be longer than FE cells and had therefore made this comparison as a one-tailed test. However, the effect observed was in the opposite direction (AE < FE), therefore *P* values are large and close to 1. Also cells recorded from subject GA had shorter latencies, on average, than those from EL. In making comparisons between subjects, we did not expect a difference, therefore permutation tests were done to test for inequality and were two-tailed (indicated with ‘<>’ in Table 2).

**Figure 3.**
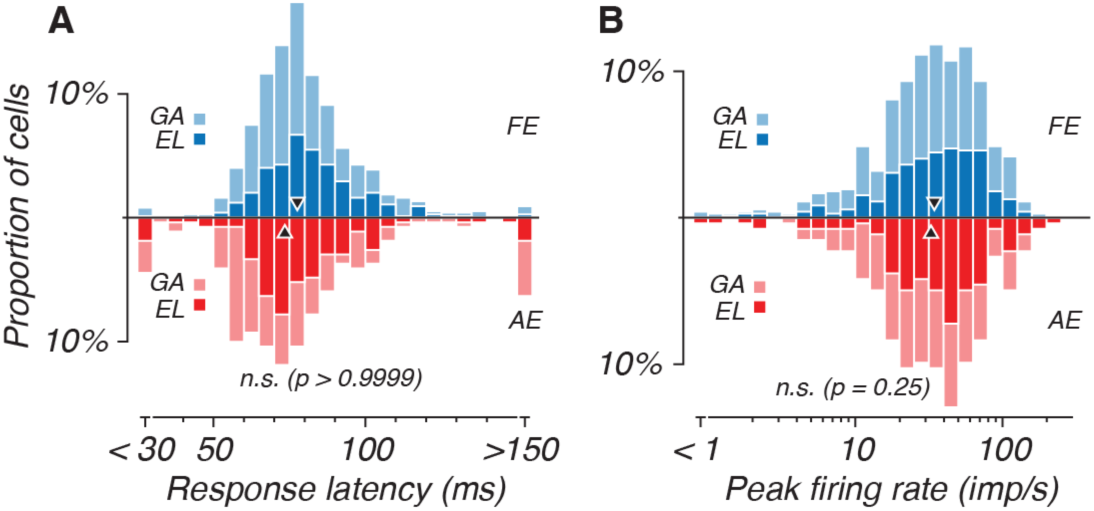
Distributions of response latency and peak firing rate for neurons from GA and EL. FE cells are colored blue, AE are red. Subject EL is dark, GA is light. **A.** Visual latency. Latencies > 150 ms or < 30 ms are grouped in single bins and considered outliers. Permutation tests with outliers excluded on GA and EL together revealed for FE: *p < 0.0001*, AE: *p < 0.01*, and testing FE > AE for GA: *p < 0.0001*, EL: *p = 0.005*, and for both subjects: *p < 0.0001*. **B.** Peak response. Mean firing rate of preferred condition. Permutation tests revealed no significant differences between subjects and viewing conditions (see Table 1).

**Table 2:**
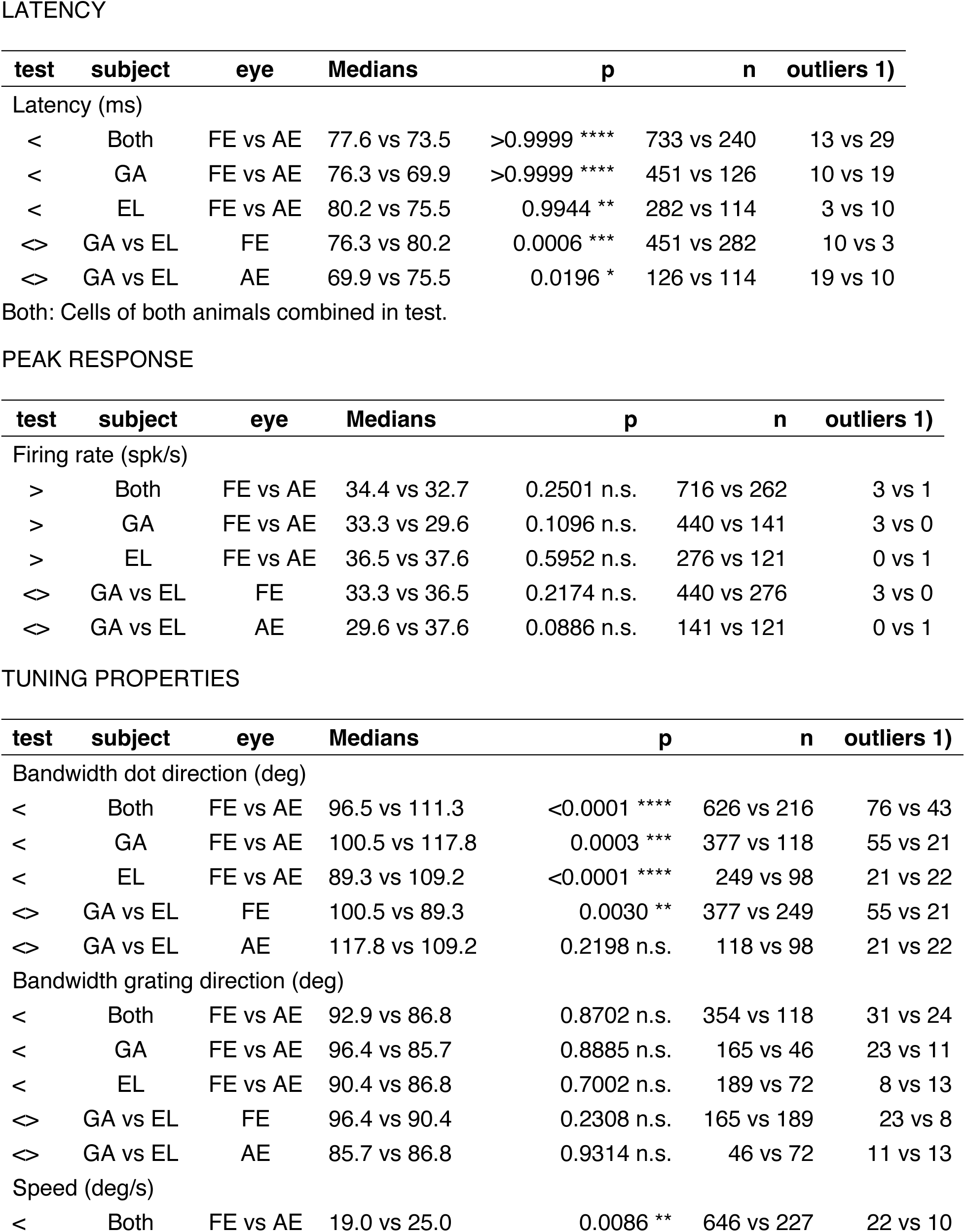
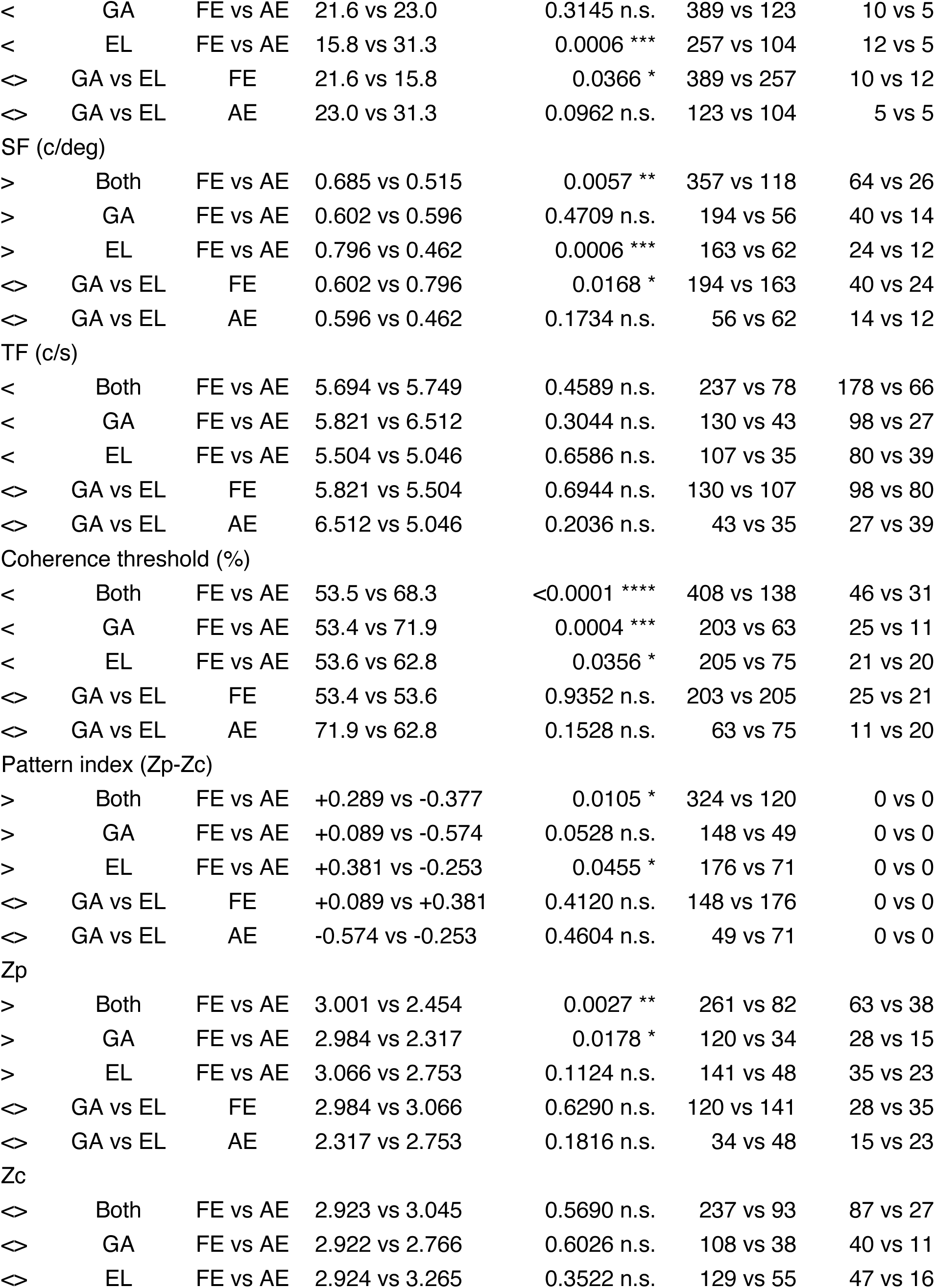
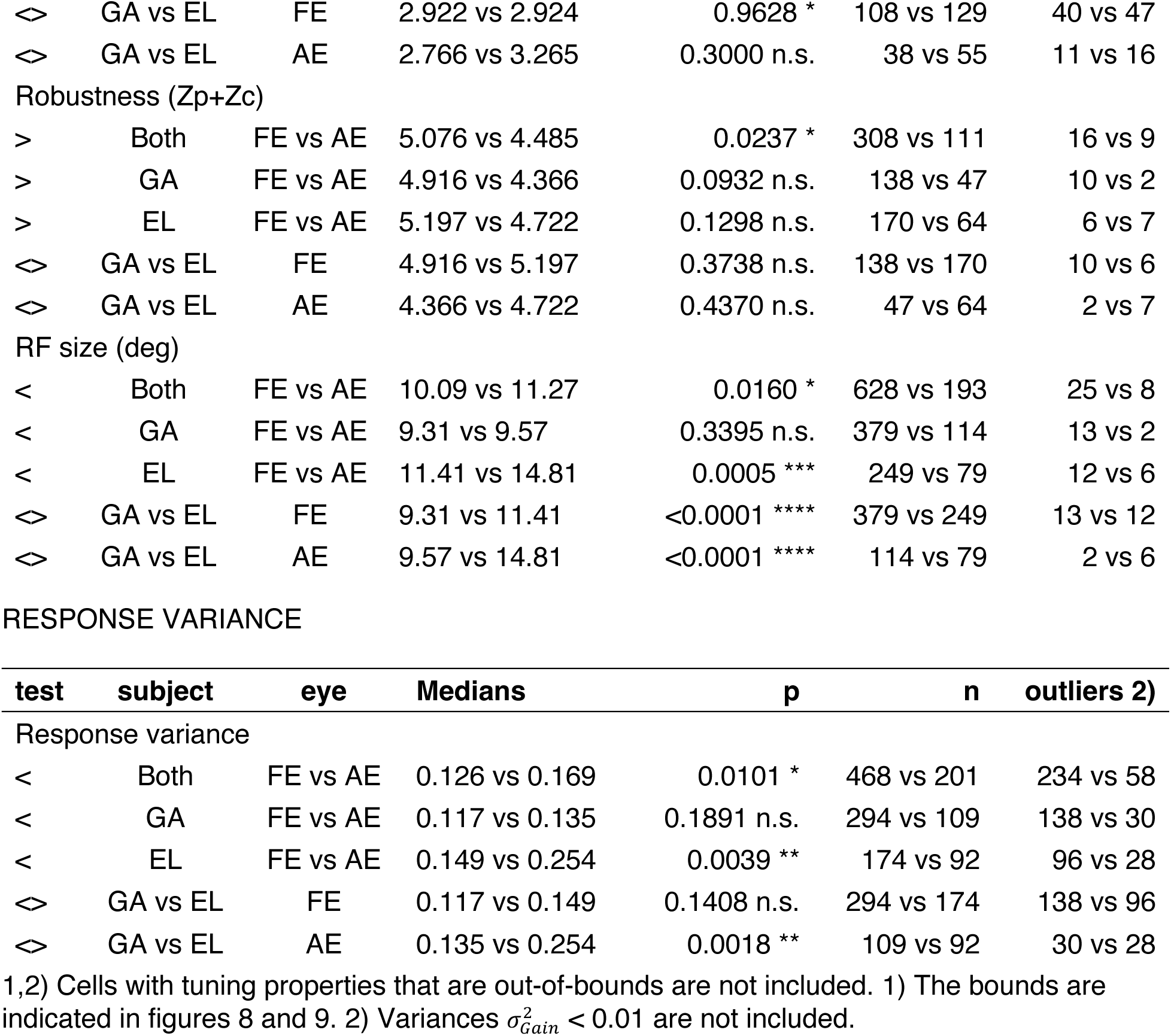
Median permutation tests.

Peak firing rate distributions are compiled in Fig. 3B. The range observed was not different from that typically seen in MT, and there were no significant differences between populations responding to FE stimulation vs. the AE or between animals (see *Table 2: Peak response*).

We routinely ran a battery of experiments on each cell to assess general tuning characteristics, including direction tuning, speed tuning, and size sensitivity. Figure 4 shows data from five example cells in two experiments: dot direction (right column) and dot speed (left column). The cells differ in their response characteristics not only by their preferred direction and speed, but also by their difference in activity recorded under FE and AE viewing. Cell 1 showed a preference for dots moving at a speed of 33 deg/s in a direction of 213 degrees (blue traces). However, when the cell was recorded under AE viewing (red traces), it did not exhibit a direction tuning profile and firing rates did not exceed spontaneous firing (left, *‘blk’*). Conversely, cell 5 was unresponsive when viewing with the FE and could only be driven through the AE. Binocular cell number 3 was tuned for ∼80 deg/s in an upward direction and its preference was maintained across eyes; activity was similar when viewing with FE and AE. Cells 2 and 4 also maintained their tuning preferences across eyes, but the activity was reduced when viewing with the non-preferred eye.

**Figure 4.**
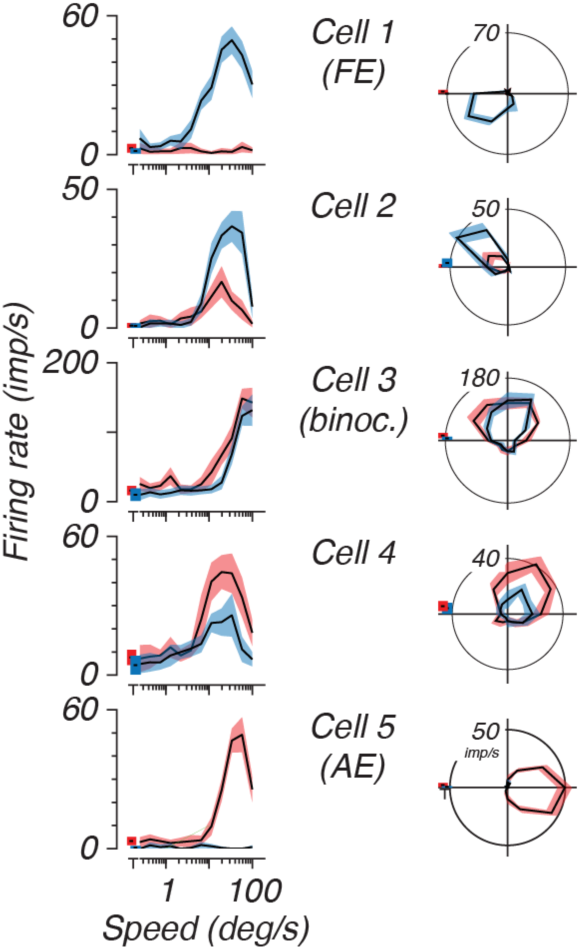
Speed and direction tuning for 5 example neurons. The left column shows speed tuning measured at the preferred direction for patches of moving randomly placed dots. The right column shows direction tuning for the same cells, in polar coordinates. Each cell’s responses measured under FE viewing (blue) and AE viewing (red) are overlaid. Mean firing rate is bounded by a shaded area representing the standard error of the firing rate. The firing rate during blank screen presentation is offset from the other conditions. Cells are ordered by eye preference from top to bottom. Cell 1 was FE-preferring and cell 5 was AE-preferring.

### Eye dominance

The range of eye dominance characteristics exemplified in Fig. 4 were observed in both animals. Unlike visually typical MT, which is largely binocular, eye dominance for these animals ranged along a continuum from exclusively FE-driven to exclusively AE-driven. Some cells even exhibited suppression in the non-preferred eye at the cells preferred stimuli. These *‘super monocular’* cells were observed among both the FE and AE populations.

Figure 5A shows classification of eye dominance for 3 example cells, one binocular (black), one purely FE responsive (blue), and one purely AE responsive (red). Figure 5B shows the distribution of ocular dominance for both animals computed as described in Methods (see equation 1). The distribution of eye dominance from these animals is strongly skewed toward the fellow eye (ODI +1). Here we calculated an ocular dominance index (ODI) from –1 to +1 by regressing firing rates of matching conditions when viewing by FE or AE (see fig. 5A).

**Figure 5.**
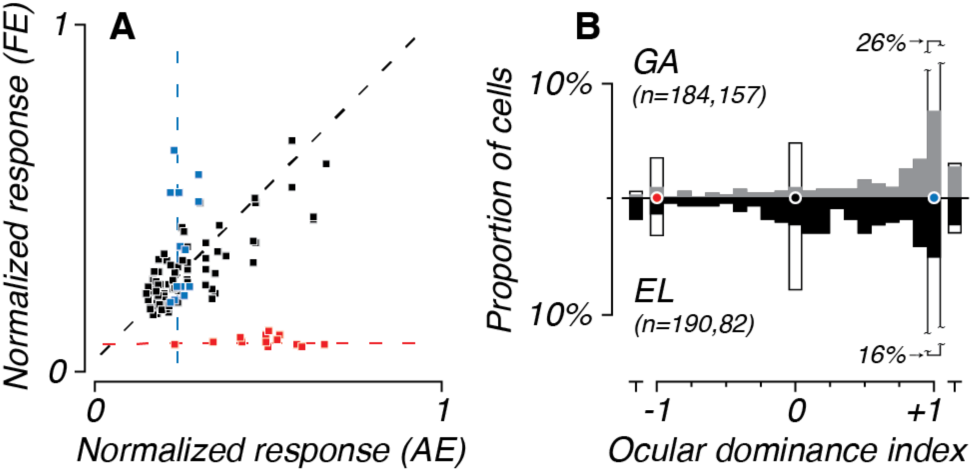
Eye dominance. **A.** The ocular dominance index (ODI) was obtained by comparing firing rates evoked in the two eyes (quantitative ODI). Responses of 3 characteristic example MT neurons during AE and FE viewing illustrate how quantitative ODIs are obtained. The firing rates obtained during FE and AE viewing is regressed for all matching stimulus conditions. Linear fits, computed taking account of variability in both directions, are indicated with dashed lines. For the case shown in black, the cell’s firing was similar regardless of the viewing eye and thus binocular. The slope is close to 1, giving an *ODI = +0.0028* (eq. 1). For the case in blue the firing rate varied when viewing with FE, but did not vary from baseline during AE viewing. The fitted line only depends on the firing rate in FE resulting in a slope close to infinity (*ODI = +1.0007*). For the case in red, the cell had constant firing during FE viewing, but varied during AE viewing. The slope of this is close to zero (*ODI = -0.9990*). **B.** Distribution of quantitative ODI values for subject EL (black, below) and GA (grey, top). The example cells from A are indicated by color-matched circles. In cases where quantitative ODI could not be done, qualitative ODI was based on the cell response during mapping experiments (open bars). A small subset of cell exhibited inhibition when stimulated through the non-preferred eye. These cells are offset at *ODI > +1* and *ODI < -1*. Some MT cells were tested only with one viewing condition and an ODI could not be obtained from these cells. These unscored cells are not included: 280 in subject GA and 154 in subject EL.

Quantitative scoring thus required that the cell was tested under both AE and FE viewing conditions. When the data did not allow matching of conditions across eyes, cells were scored qualitatively (open bars) by assessing audible spikes. ODI measures were not obtained from cells that were tested through only one eye and are not included in the bar graph (280 in Subject GA, 154 in subject EL). Cells that showed suppression in the non-preferred eye are in bins < –1 and > +1.

MT cells in normally-reared animals are almost always binocular (Maunsell & Van Essen, 1983), as they were in our recordings from the anesthetized control animal (Fig. 11A). Figure 5 shows that there was a limited number of binocular MT cells in the amblyopic animals, and that AE preferring cells were underrepresented. When comparing subjects, GA’s distribution showed a stronger shift toward monocular FE cells, while EL, although biased towards FE, had a stronger representation of the AE.

### Motion coherence sensitivity

A striking behavioral deficit in amblyopia is reduced sensitivity for motion coherence (Fig 1C and D). It is therefore of interest to evaluate the coherence response properties of MT neurons in these amblyopic animals. Figure 6A shows coherence-response functions for two example neurons. As is typical in MT, firing rate increases with increasing dot coherence, with apparent motion in the cell’s preferred direction and speed. Both cells show an increase in firing rate as the coherence increases, although the FE cell (blue) seems to rise at a faster rate than the AE- preferring cell (red). The rate at which the firing rate increases as a function of coherence reflects the cells coherence sensitivity.

**Figure 6.**
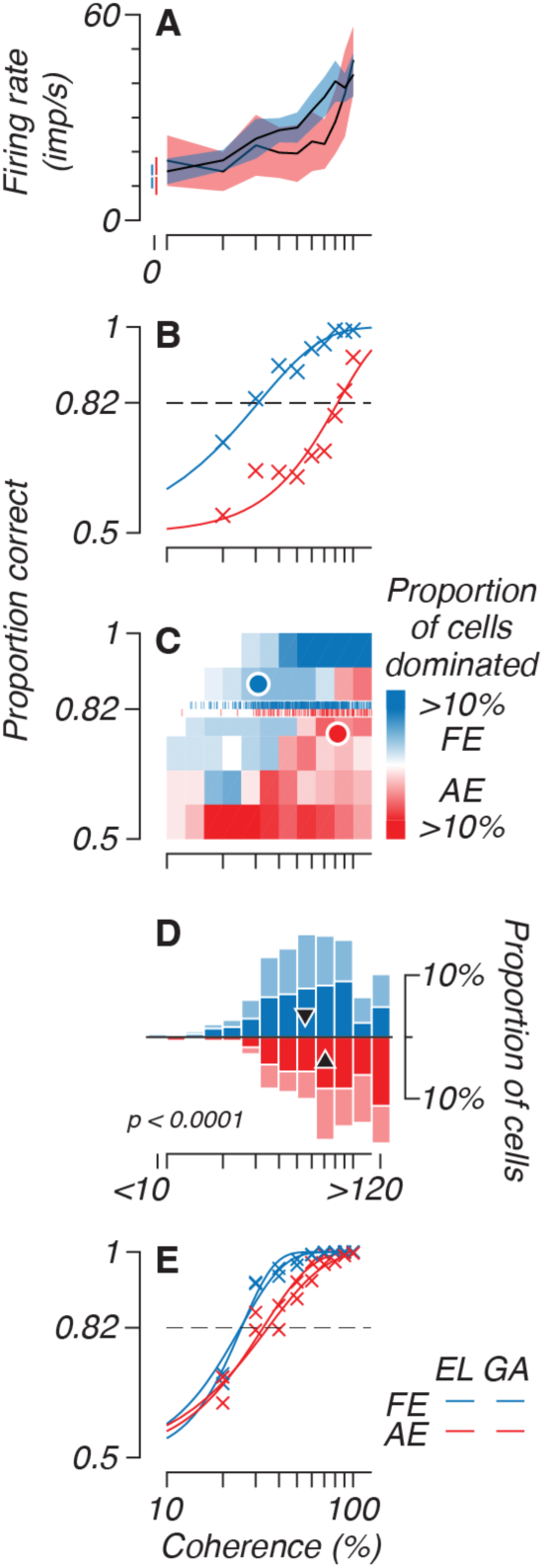
Response to motion coherence. **A.** Firing rates of two example cells from subject GA. Shaded areas indicate standard deviation around mean firing rates. In blue is a cell that preferred the FE and in red a cell that preferred the AE. Firing rates obtained during random noise (0% coherence) is off-set (left) from the conditions with non-zero coherence. **B.** Detection performance of the same two cells as panel A. Performance is obtained by calculating the surface of an ROC when comparing coherence trials with trials of 0% coherence. Performances are fitted with a Weibull curve. (Note that curves are extrapolated past 100% coherence. For details see Methods). Cell’s thresholds are indicated at the intercept of the fitted curve with a performance of 0.82 (Threshold FE is 31% coherence and 83% for the AE) **C.** Detection performances of all cells. Thresholds are indicated with individual vertical line segments (rug plot) at a performance of 0.82. The thresholds of the example cells in panels A and B are indicated with color-matched circles. The underlying color code indicate the overrepresentation of FE (blue) or AE (red) fitted curves within a rectangle bin. Curves running through higher coherence bins (left) are mostly FE cells. **D.** Distribution of coherence thresholds. Medians are indicated with triangles. The median threshold for the FE was significantly smaller than for the AE (*p < 0.0001*). **E.** Threshold computed for a simulated pool 10 of cells (details in Methods).

We used the method described by Britten et al. (1992) and El-Shamayleh et al. (2010) to obtain coherence thresholds for each cell (see Methods). We compared the spike counts of 0% coherence conditions (isolated bars at the left in Fig. 6A) with the spike counts of each condition with coherence >0%. This function also represents the ability of a cell to detect motion, and can therefore be expressed as a performance measure (Fig. 6, panels B, C and E). Figure 6B shows the performance of the same two cells as in panel A. The intercept with 82% is taken as the cell’s coherence threshold (see Methods). Of the two example cells, the threshold of FE preferring cell is lower than the AE preferring cell.

Figure 6C shows the percentage of cells dominated by each eye at the corresponding coherence (abscissa) and performance (ordinate) level. Blue rectangles represent FE- dominated cells, red are dominated by the AE, and equal influence is white. It is apparent that the majority of cells with low coherence thresholds are dominated by FE and those with high coherence thresholds are dominated by the AE. The small vertical markers at 82% represent the thresholds of all cells recorded. Figure 6D shows the distribution of these thresholds. The distributions do not differ between the animals (FE: *p = 0.91*, AE: *p = 0.15*), but overall coherence threshold is lower for FE cells (GA: *p = 0.001*, EL *p = 0.03*, both subjects: *p < 0.0001*).

Since psychophysical performance is based on the combined activity of many neurons, we performed a pooling analysis similar to that described in El-Shamayleh et al. (2010) (see Methods also) to simulate the performance of the population. Figure 6E shows the neurometric functions obtained with this method using a pool size of 10 neurons (changing the pool size does not alter the conclusion, as shown in supporting figure S1). As with the single neuron analysis, the threshold is lower for the FE when neuronal activity is pooled. In this case, we used equal pool sizes for the AE and FE, but in reality FE-dominated neurons outnumber AE cells (figure 3). The difference in threshold between the two eyes’ populations would be even larger if we used pool sizes based on the actual ratio of FE to AE cells (cf. Shooner et al., 2017).

### Pattern motion selectivity

Pattern selectivity can be examined by presenting two superimposed gratings moving in different directions, which yields a plaid pattern when fused into a coherent moving image. For identical gratings separated by 120 degrees and drifting in their respective directions, the motion direction of the plaid is midway between the directions of the two components (e.g. 0 deg with components moving at -60 deg and +60 deg). By measuring the motion response to a single grating, the response to plaids can be predicted. If a cell is selective for the pattern direction, the response typically shows one peak, at the same angle as for a single grating. However, when a cell’s response is determined by the individual components of the plaid, the response peaks when one of the components is drifting in the preferred direction, resulting in a response with two peaks (Movshon et al., 1985; Rust et al., 2006).

The integration of motion components into a global representation is an advanced form of encoding and continues to develop after birth (Kiorpes and Movshon, 2014). It is therefore of interest to see whether amblyopes, who experience disordered visual experience during development, show different pattern selectivity in cells that respond to the AE and FE.

We classified cells as component or pattern-selective by comparing their responses to plaid stimuli against two predictions generated from the response to gratings (Movshon et al.,1985; Smith et al., 2005). This results in two Z-transformed correlation coefficients: one for the pattern model (Z_p_) and one for the component model (Z_c_). A pattern selectivity index is computed by taking *PI = Z_p_ – Z_c_*. For pattern-selective cells, PI > 0. For component-selective cells, *PI < 0*. Significance levels are *Z* ± 1.28 (see Methods). Cells that do not reach significance are considered *unclassed*.

Figure 7 shows a scatter plot of pattern (Z_p_) and component directional (Z_c_) selectivity. Component selective cells are in the lower-right of the plot (high Z_c_ and low Z_p_) and pattern selective cells are in the upper-left. The diagonal histogram on the right shows the distribution of the pattern index for FE (blue) and AE (red) preferring cells. The histogram on the left shows a distribution of ‘robustness’ of the Z-scores, a measure of the overall statistical reliability of the response. The open triangles indicate median values. Both pattern index and correlation robustness are significantly different between the eyes. Inspection of the mean values of Z_p_ or Z_c_ for these populations (indicated with plus markers) shows that the difference is entirely in the Z_p_ direction. The difference between the eyes can be explained by a diminished pattern correlation in the AE. Table 2 (Tuning properties, pattern index) shows that the differences between the eyes are not significant in Z_c_ but are significant in Z_p_.

**Figure 7.**
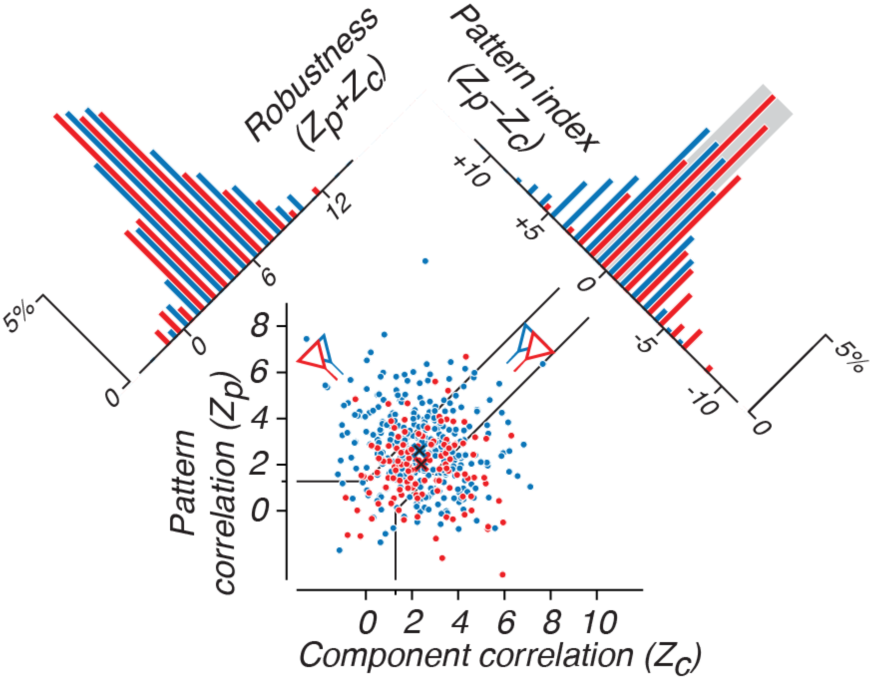
Pattern selectivity. Responses to plaid stimuli can be tuned to the global pattern direction or along the underlying component directions. Cell’s grating responses are used to predict idealized pattern or component responses to the plaid stimulus. Correlation with pattern selective model is Z-transformed and expressed as coefficient *Z_p_*. *Z_c_* indicates the component coefficient. Cells above the unity line correlate better with a pattern direction response. Component selective cells are below. The differences between *Z_c_* and *Z_p_* reflect pattern selectivity of a cell and is expressed as the pattern index: *PI = Z_p_ – Z_c_*. The differences between *Z_p_* and *Z_c_* are not significant for correlations below *1.28* (for *Z = 1.28*, *p = 0.9* indicated by solid lines). Both cells preferring FE (blue) as cells preferring AE (red) have pattern and component selective cells. The distribution of pattern indices is shown in the right histogram orthogonal to the unity line. Note that pattern selective cells (*PI > 0*) are shown up and left. Permutation test reveal that the pattern selectivity is larger for FE when subjects are pooled (*PI = +0.289 > -0.377*, *p=0.0090*). This is also significant for subject EL (*PI = +0.381 > –0.253*, *p=0.0461*), but not for subject GA (*PI = +0.089 > –0.574*, *p=0.0587*). Differences between subjects are not significant (two-tailed, FE: *p=0.4180*, AE: *p=0.4782*). Robustness indicates a strong correlation of the pattern response to either prediction based on the single grating response and is the sum of the correlation coefficients: *Z_p_ + Z_c_*. The robustness of the responses are shown in the left histogram, with plus signs indicating the median values for both the FE and AE. These indicate higher robustness for FE. This is only (barely) significant when subjects are pooled (*Robustness = 5.076 > 4.485*, *p=0.0238*), suggesting that the differences in response between the eyes are modest. For details see Table 1.

Thus, although some AE cells still showed pattern selectivity, this was reduced overall. This could in principle result from changes in either V1 or MT, an issue we will take up in the Discussion.

### Interocular differences in motion selectivity

We have until now considered single-neuron measures of response and sensitivity. But differences in the coding properties of neural *populations* might more accurately represent the information available for behavior, and account for the interocular difference in motion coherence sensitivity. Neural populations for the AE and FE may be tuned to different stimulus ranges or be more or less selective. We looked at tuning bandwidth, range – or selectivity – and dot speed and coherence, as well as grating tuning properties (see figure 8).

**Figure 8.**
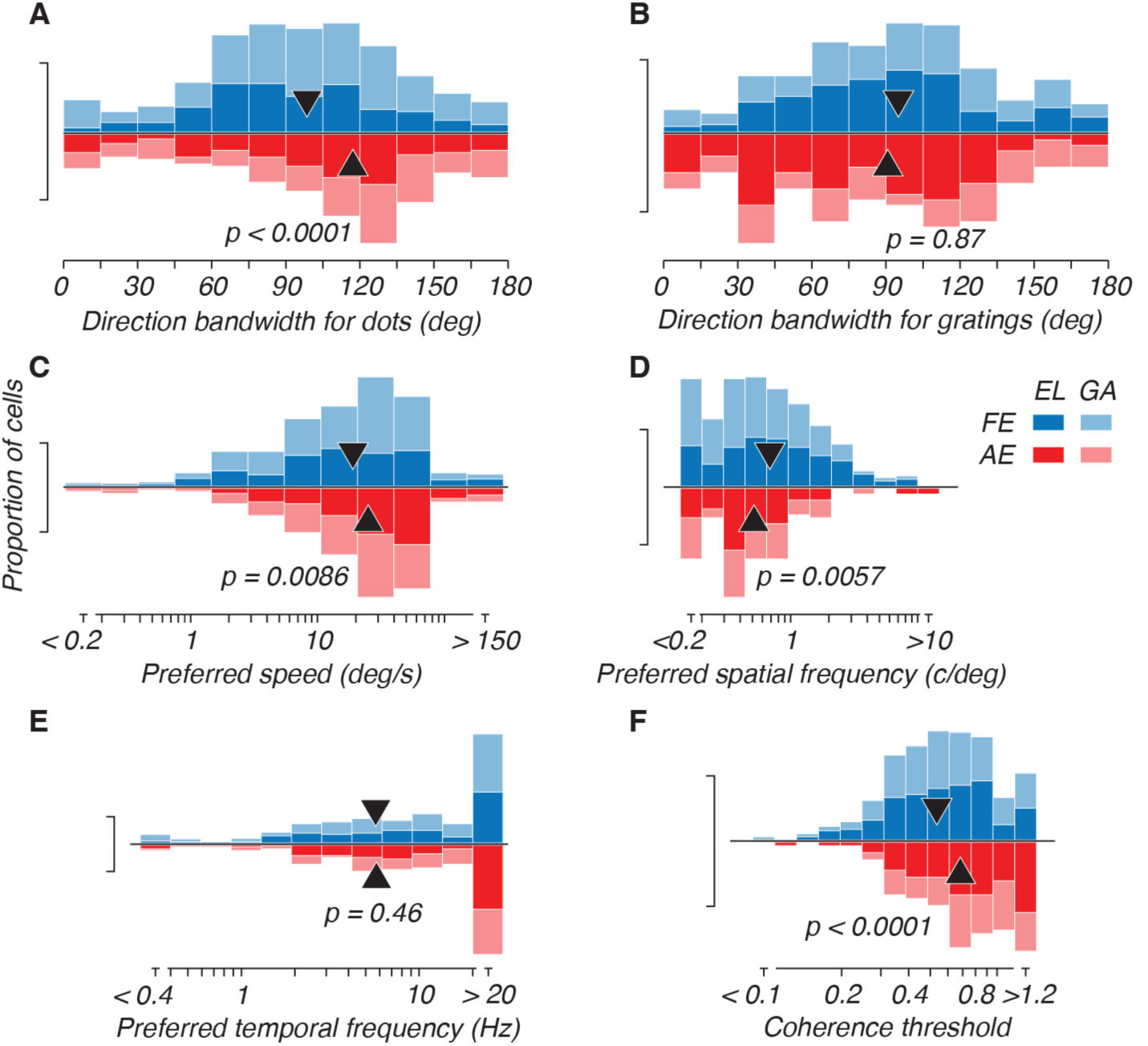
Population differences of tuning properties. Distributions of tuning are shown for FE blue (top) and red (below) AE. Dark colored bars represent subject EL and light colored bars subject EL. The whiskers indicate 10% of the total number of tested cells. The p-values are the result of permutation tests between eyes with subjects pooled (black triangles represent the medians of those pooled distributions). Bins with a greater-than or less-than sign (*>* or *<*) indicate outliers and are excluded from permutation tests. The tuning distributions are presented for **A.** Bandwidth of dot direction; **B.** Bandwidth of grating direction; **C.** Preferred speed for dots; **D.** Preferred spatial frequency of gratings; **E.** Preferred temporal frequency of gratings; **F.** Dot coherence threshold for optimal conditions. The direction bandwidth difference is significant for dots (smaller in FE: *p<0.0001*), but not significant for gratings. TF tuning was not significantly different between eyes. Detailed statistics can be found in table 1.

For moving dots, we found that tuning bandwidth was narrower for FE than for AE cells when pooled (*p < 0.0001*, figure 8A, *Table 2: Tuning properties*). This difference is significant for the subjects individually as well (GA: *p = 0.0003* and EL: *p < 0.0001*). Comparing across subjects, the FE pool is on average more narrowly tuned for EL than for GA (*p = 0.0030*), but there was no significant difference between subjects for the AE (*p = 0.2198*). Direction tuning bandwidth differences were not significant for grating stimuli.

Consistent with behavioral findings, the median preferred speed of the FE population was slower than for the AE when subjects are pooled (*p = 0.0086*) and for EL alone as well (*p = 0.0006*). While the pattern was the same for GA, the difference in preferred speed between eyes was not significant. Speed tuning was different between subjects for the FE (*p = 0.0366*, GA median was slower); there was no inter-subject difference for the AE.

The results for spatial frequency tuning are consistent with those for speed tuning. See Table 2 for statistics. All the population differences for temporal frequency were non-significant. This indicates that the difference in speed sensitivity is mainly due to a shift in SF sensitivity alone.

### Response variance

The poor visual sensitivity of the AE might result from reduced reliability of neural responses or higher response variance (El-Shamayleh et al., 2010; Wang et al., 2017). A highly reliable neuron would respond nearly identically to every identical visual stimulation. Typically, this quantity is expressed as a Fano factor (the ratio of spike count variance to mean). However, Goris et al. (2014) made an improved model of response variance by introducing a gain factor (variance of gain: +^-^ , eq. 2) whose impact depends on response magnitude. We fit this model for every cell and present the gain factors in Figure 9. As expected, on average the variance increases with mean firing rate (Fig. 9A) but in this case non-linearly, indicating a gain factor adding extra response variance (variance of gain). The curved lines are median model fits bounded by 95% for FE (blue) and AE (red). A Fano factor model would have yielded straight lines parallel to the diagonal and would have been a poorer description of the cells’ response.

**Figure 9.**
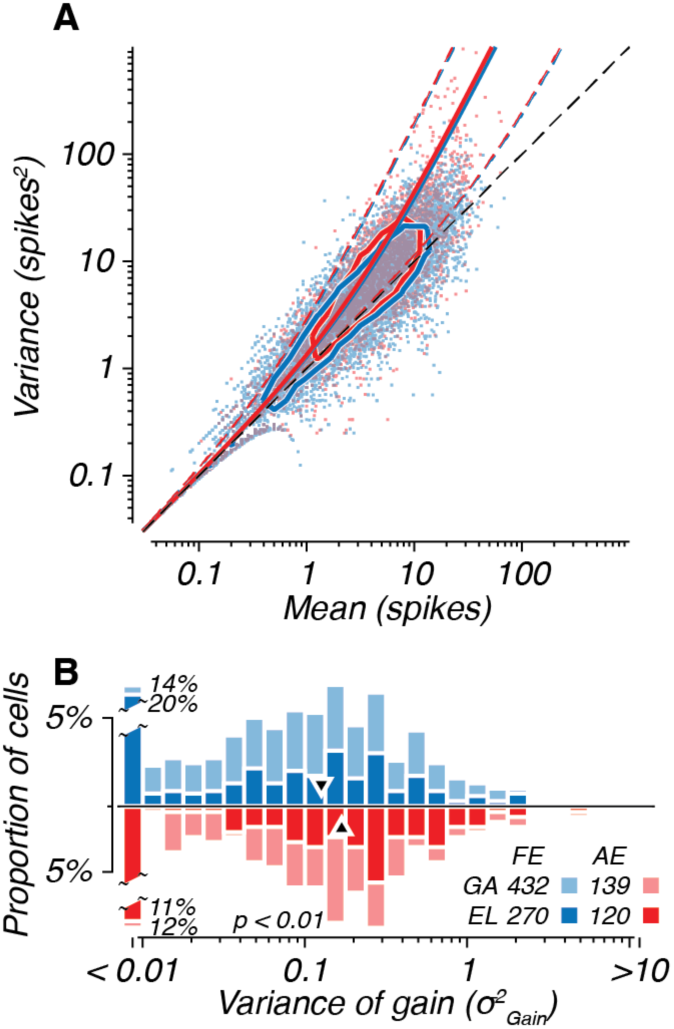
Response variance. **A.** Mean firing and its variance is computed for all conditions in direction tuning experiments. Oval shapes are half-max densities of the underlaying data clouds for FE (blue) and AE (red). Solid curves indicate the mean fit values of the *variance of gain* 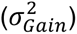 of all cells. Dashed curves bound 95% of the observed values. The unity line indicates a Fano factor of 1. Different Fano factors values would run parallel to this unity line. Note that a repetitive pattern can be seen at low mean spike counts (<1 spikes), this is due to the discreteness of counting spikes within a stimulus epoch. **B.** Histogram of fitted values of 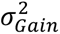. The difference between the eyes (pooled for both subjects) is significant (Permutation test with outliers excluded: *p<0.01*), with the median predicted variance of gain being higher for the AE, suggesting less reliable firing when viewing through the AE. Detailed statistics can be found in Table 1.

Interestingly, 34% of FE and 23% of AE cells exhibit virtually no modulation with increased response rate (+^-^ < 0.01), which is also captured by these fits and occurs when Fano factors are close to 1. Any difference between FE and AE cells does not appear to be large: data points and the fitted lines largely overlap. However, the small difference in variance of gain is significant between the eyes (*p = 0.0101*); with FE having a lower factor indicating higher reliability. A significant difference is maintained for subject EL alone (*p = 0.0039*), but not for GA alone (*p = 0.1891*). Reliability is different between subjects for the AE (*p = 0.0018*), with GA’s AE showing more reliable responses than EL’s AE, which is interesting given GA’s less stable fixation. Because gain variance relates to state changes with a relatively slow time course (Goris et al., 2014), the lower values for the FE could reflect a more consistent pattern of attentional deployment when using the FE. It might also reflect the less consistent patterns of fixation seen in both animals when using their AEs. Table 2 summarizes these results (Response variance).

### Interocular tuning correlations

The majority of the cells responded well only to visual stimulation of one eye. However, there was a small subset of binocular cells with which we were able to compare responses under both viewing conditions for a single cell. To be included in these comparisons, the cell needed to 1) be recorded under both FE and AE viewing condition, and 2) have a reliable response when each eye was viewing. We constrained our analyses to binocular cells with an eye dominance between *-0.25 < ODI < +0.25*.

In Figure 10, tuning properties are compared between the eyes for cells responding under both FE and AE viewing conditions. Cells with close-to-binocular ODIs (*ODI ≈ 0*, black data points) seem to cluster around the diagonal, whereas light-grey (monocular) cells lie further from the diagonal. Estimates are based on the stimulus conditions that evoke close to maximum firing rates in either eye, and thus will be less influenced by noise. Less reliable estimates through one eye could obscure similar tuning and as a result give rise to offsets in cells that are not balanced across eyes.

**Figure 10.**
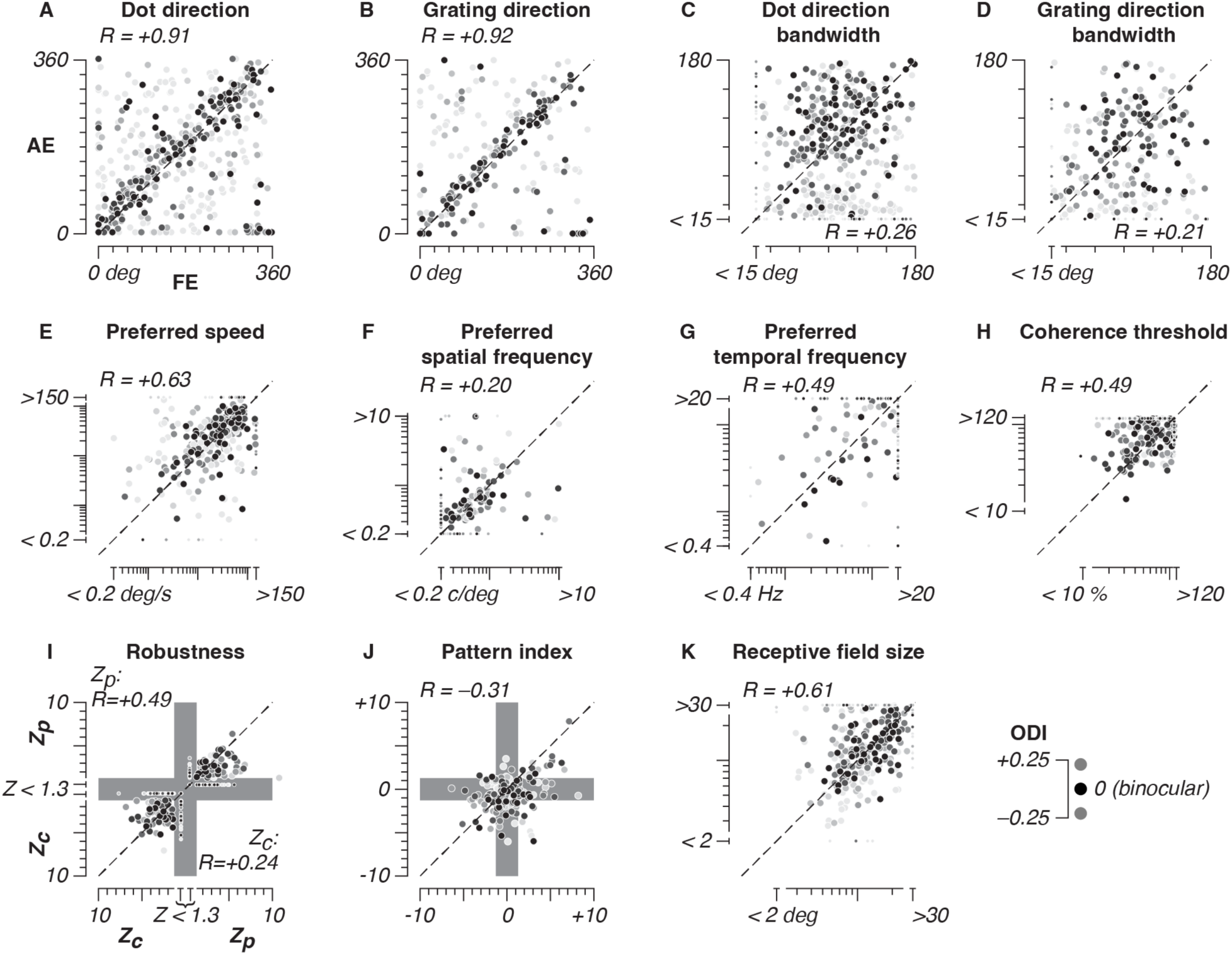
Interocular correlation of tuning. Binocular cells are indicated in black. Other cells are grey-scaled according to the ODI. Pearson’s correlation coefficients are shown for values obtained during FE versus AE viewing. For the regression analysis monocular cells (*-0.25 > ODI > +0.25*) are excluded, as are cells that have tuning values that are out of bounds. Outlier values are shown as small symbols around the margin of the panels. Correlation was performed after transformation to logarithmic scales for panels E-H and K. Direction bandwidth tuning difference exceeding 180 deg are corrected to avoid wrapping around 180 degrees (Panels C and D). Tuning is presented for **A.** Dot preferred direction; **B.** Grating preferred direction; **C.** Bandwidth of dot direction tuning; **D.** Bandwidth of grating direction tuning; **E.** Preferred speed for dots; **F.** Preferred spatial frequency; **G.** Preferred temporal frequency; **H.** Dot coherence threshold; **I.** Correlation coefficients for predicted pattern direction response (Z_p_) and component direction (Z_c_). The gray shading indicates non-significant correlation coefficients (see also fig. 7). **J.** Pattern index. Significance bound also indicated in gray; **K.** Receptive field size. The two axes of the fitted two-dimensional Gaussian were averaged.

Pearson’s correlation coefficients are indicated inside each of the panels in Figure 10. Tuning direction was similar between the eyes (Fig. 10A and 10B, *r = 0.91* and *r = 0.92*). Both for dots and gratings there are no systematic deviations from the diagonal. However, neither grating nor dot direction bandwidth correlate well and dot direction bandwidth appears broader for AE since data is above the unity line, consistent with results from the population as a whole (Figure 8).

Despite substantial behavioral deficits, and the population differences noted above, dot speed tuning did not differ for binocular cells when recorded under FE or AE viewing (*r = 0.63*).

Moreover there was no systematic shift toward faster preferred speeds for the AE in this population, unlike for the population as a whole. This suggests that the tuning properties of binocular neurons remain better matched than the overall population properties of the monocularly driven neurons with which they are intermingled.

As noted earlier, (Figure 2B), the stimulus locations had to be shifted systematically across eyes to achieve peak firing rates, but the size of the receptive fields matched for the two eyes (Fig. 10K, *r* = 0.61) with no systematic shift to a larger size for AE.

### Recordings under anesthesia

After completing the awake behaving experiments, we made additional recordings in subjects EL and GA and in a third age-matched control (subject PF) under opiate anesthesia. We aimed to compare the awake amblyope recordings directly to measurements made under anesthetized conditions, which we described previously for a different set of animals (El Shamayleh et al. 2010). In addition, we took advantage of the greater control over optics and eye movements afforded by the anesthetized preparation to determine whether and how the fixation instability that we had observed (Figure 2) influenced our results.

The responses of neurons were recorded in area MT and the surrounding extrastriate areas with linear probes in response to three sets of stimuli: sinusoidal gratings varied in orientation and spatial frequency; random dot fields varied in speed and direction; and sparse noise. We derived tuning measures from these stimuli to compare to the awake-behaving recordings, and used them to map the spatial extent of receptive fields of many neurons recorded simultaneously.

The statistics of our neural data recorded from area MT under anesthesia aligned well with those made in the animals when they were awake and fixating. Figure 11A illustrates the distribution of ocular dominance indices for our three animals. The data distribution from GA and EL aligned well with the data recorded in the awake preparation. The only substantial difference between awake and anesthetized measurements was in EL. For EL, many recording sites showed weak excitation to amblyopic eye stimulation, but none showed the dominant amblyopic eye input that was observed in both subjects in the awake preparation and in GA in the anesthetized preparation. Two aspects of this animal’s recordings could explain this difference. One is the low number of neurons recorded (N=53) in EL, and the second is that most recorded sites were in areas V3 and V4t on the upper bank of the STS rather than area MT (see methods for further details). We believe that the low number of neurons recorded is most likely, since both areas V3 and V4t are binocular under normal visual development, and the bias towards fellow eye representation is still consistent with a broad reorganization of the binocular selectivity of extrastriate cortex in amblyopia. Indeed, both subjects’ distributions are markedly different from the distribution in subject PF, our age-matched normal control.

**Figure 11.**
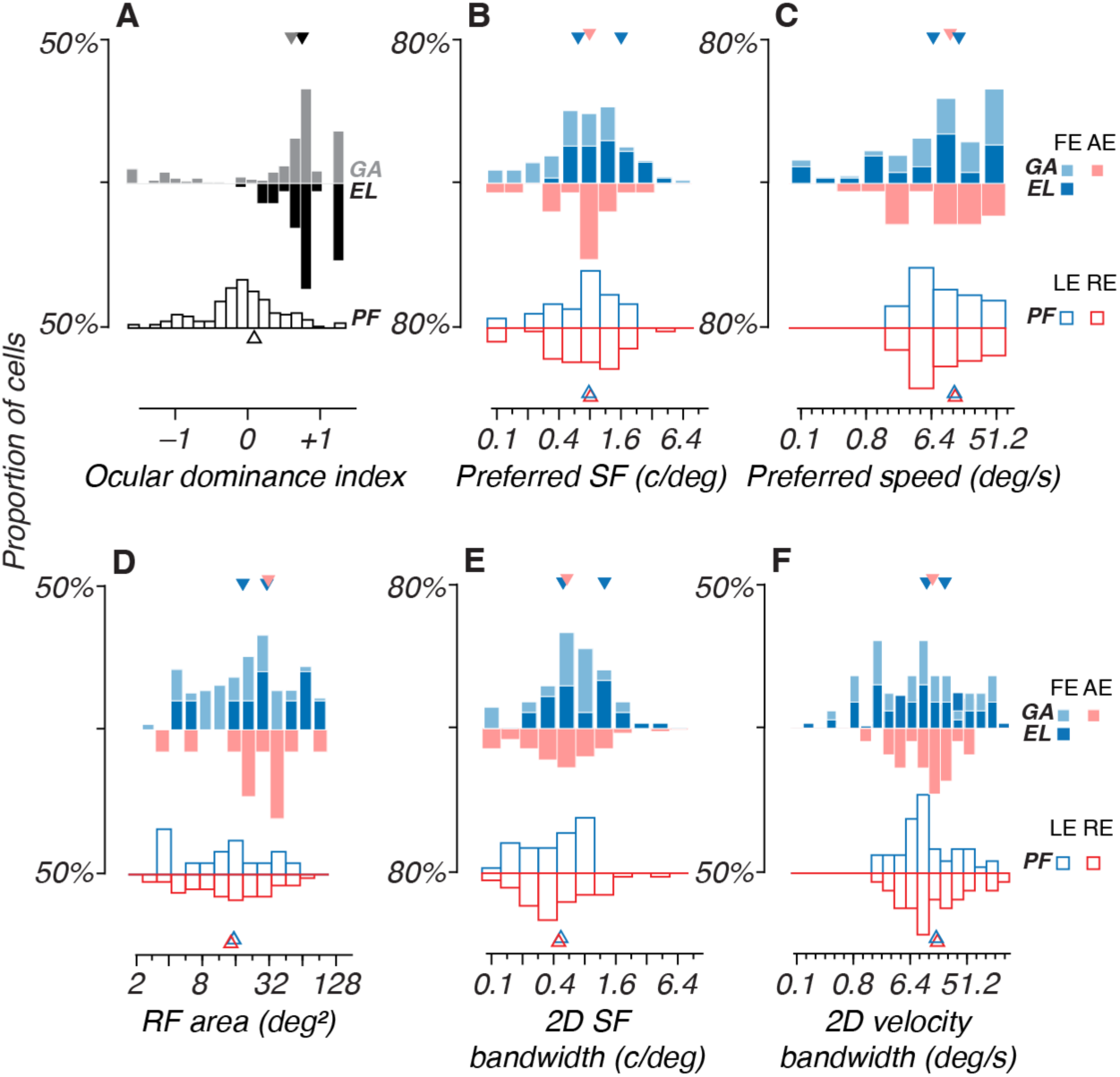
Neuronal selectivities in anesthetized macaques. In each plot the distribution of properties of amblyopic subjects is shown above as a north-south pair of histograms with the fellow eyes on the top and the amblyopic eyes on the bottom. Data for the age-matched control subject with normal vision (PF) are given at the bottom of each subplot in similar format. **A**. Ocular dominance indices for each animal. Gray bars show subject GA and black bars show subject EL (as in Figure 3). Unfilled bars show the distribution of ocular dominance indices for the control animal, comparing left eye to right eye responses. The triangles indicate the mean of the cells with ocular dominance indices between –1 and +1. Subplots **B-F** follow the same color conventions as Figure 3. Triangles indicate the means of each distribution. **B, E**. Spatial frequency preference and 2D spatial frequency bandwidth of fellow and amblyopic eye-biased cells in each subject. **C, F**. Distributions of speed preference and 2D velocity bandwidth (square-rooted). **D**. Receptive field area. The receptive field area is the product of the diameter of each cell’s receptive field estimated in response to moving dots and fit using the function described in the Methods section.

Figure 11 B, E and D, F illustrate the preferred spatial frequency and speed in the fellow and amblyopic eye in subjects GA and EL, along with the bandwidths for spatial frequency and speed. Because the stimuli varied in both direction and speed/spatial frequency conjointly, bandwidth is shown as the square root of the two-dimensional product of speed and direction bandwidth (hereafter, “2D velocity tuning”). The preferences for spatial frequency and speed and the slight differences in preference between eyes were well aligned with the awake data presented in Figure 8. In subject GA, only 2D velocity bandwidth differed between eyes (16 vs 26 deg/s^2^, p=0.03 Kolmogorov-Smirnoff (2-sample test). All other interocular differences failed to reach statistical significance. In fact, they were all smaller than the magnitude of differences between subjects EL and GA in their fellow eyes. Compared to subject GA, fellow eye cells in Subject EL preferred higher spatial frequencies (p=0.003, 2 sample KS test), and had larger 2D bandwidth (p=0.0018, 2 sample KS test). Speed preference and speed bandwidth were marginally larger in EL than in GA, but this difference eluded significance (p=0.054, 2 sample KS test). The differences in receptive field area were also unremarkable between eyes and across animals (see Figure 11D). The limited sample size in EL compared with GA made it difficult to determine whether differences between animals were a consequence of the degree of amblyopic deficit observed in each animal. However, it is suggestive that subject EL’s behavioral contrast sensitivity had a higher spatial frequency cutoff than subject GA and this is paralleled by the spatial frequency preference of sampled neurons in the extrastriate cortex.

## Discussion

This study is the first characterization of extrastriate cortical function in awake behaving amblyopic macaques. Our data, based on recordings under fixation control in area MT, demonstrate neural correlates of behaviorally measured amblyopia. In particular, in addition to a surprisingly dramatic under-representation of the amblyopic eye, we found both reduced coherence sensitivity and a shift in the range of preferred speed to higher speeds for the amblyopic eye compared with the fellow eye, which is the predicted pattern based on the behaviorally measured amblyopic deficit. Importantly, the results of this study confirmed and extended the findings of our previous characterization of neural responses in area MT of amblyopic macaques recorded under anesthesia (El-Shamayleh et al., 2010). Moreover, we ultimately also recorded from MT in the same animals under anesthesia and confirmed the basic findings of our awake recordings. This is significant because it demonstrates that identified neural deficits were not due to abnormal fixation or oculomotor control.

### Eye dominance

A striking result of our recordings is the dramatic impact on binocularity that was evident in the amblyopic monkeys. More than 90% of neurons in normal MT are binocular, as exemplified by our control animal in Fig. 11A. In the amblyopic animals, the distribution of cells in MT was distinctly monocular with the AE being strongly underrepresented (Fig. 5; 11A). Our ocular dominance distributions were heavily skewed towards monocular FE cells, despite the fact that our sampling procedure was designed to maximize the yield of AE neurons. Eye dominance observed in the same animals under anesthesia was also dramatically skewed. This loss of binocularity and skewed ocular dominance is consistent with that reported by El-Shamayleh et al. (2010) in amblyopic MT, but it is in clear contrast with observations in V1 wherein amblyopes’ eye dominance is similarly monocular but the cell counts are usually more balanced between FE and AE neurons (Movshon et al., 1987; Kiorpes et al., 1998; Bi et al., 2011). MT’s most important input projections are from cortical area V1 (Born and Bradley, 2005; Felleman and Van Essen, 1991; Movshon and Newsome, 1996). These results show that the imbalance we see in MT is not simply inherited from V1, and indicates *de novo* deficits arising at the level of MT. It is therefore also possible that additional deficits appear in areas further downstream, that may not be captured yet at the level of MT. Indeed, Simmers et al. (2006) show an amblyopic deficit in processing of optic flow patterns, which is likely to be subserved at the level of MSTd.

Since MST receives much of its input from area MT, the disrupted pattern we identified may provide evidence for this kind of cascade of impact.

### Response magnitude

As noted above, we found the pattern of AE responses in MT to be markedly different from observations in V1. Whereas in V1 most neurons are responsive to AE input in amblyopic animals, at least to some degree, we noted that activity in MT under AE viewing was often not responsive or responsive but not tuned. This resulted in eye dominance distributions that were mostly monocular and skewed toward the FE. A potential confound for the observed absence of a tuned response is that the AE cells could have lower spike counts. Identifying clear tuning characteristics might depend on peak firing rate. However, we found no difference in peak response rate between AE and FE populations (Fig. 3B). Also in anesthetized amblyopes there appears to be no difference in peak firing rates (El-Shamayleh et al., 2010). However, in figure 3 we only analyzed peak firing rate per cell. There might still be a difference for certain stimulus types.

To directly address this we correlated our measured tuning values with response magnitude. Figure 12, Table 3 and supplemental figure S2 show a more detailed analysis of the dependence of response magnitude on tuning parameters. It reveals that in fact several tuning measures were related to response magnitude. A positive correlation indicates that the value of the tuning property (e.g. bandwidth) increases with response magnitude. Conversely a negative correlation indicates that the tuning property is lower with larger responses. For instance, on average, dot direction bandwidth estimates correlate with firing rate (*R≈+0.3*). However, preferred spatial frequency was inversely correlated with firing rate (*R<0*). The direction and magnitude of the correlations were similar for both eyes and subjects for each tuning property we examined. Therefore, we conclude that the pattern of our data is not likely to be impacted by any subtle differences in firing rates between AE and FE populations.

**Figure 12.**
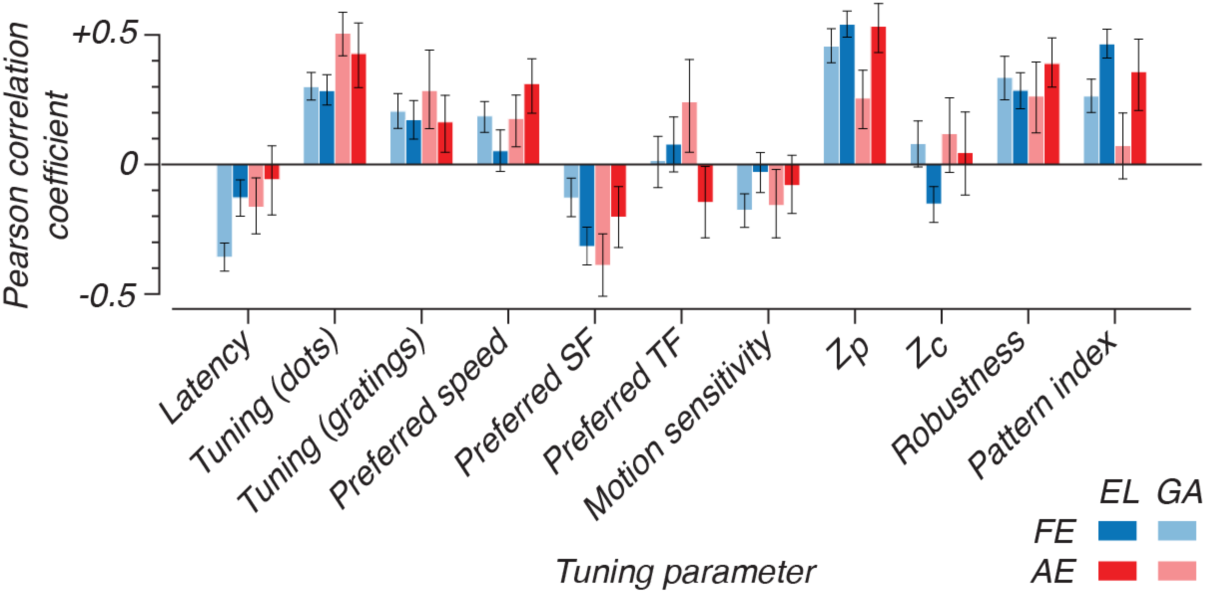
Dependence of response magnitude on tuning and sensitivity. Pearson’s correlation coefficients are shown for correlations between the log of peak response and 11 tuning parameters. Standard deviations obtained by bootstrapping are indicated (n=1000). Blue values are tuning parameters recorded under FE viewing and AE values are in red. Subject EL is indicated with dark colored bars and subject GA with light-colored bars. Speed, spatial frequency, temporal frequency, latency and coherence were log-transformed before computing correlations. The data supporting this summary are shown in supplemental figure S2 and Table 3.

**Table 3:**
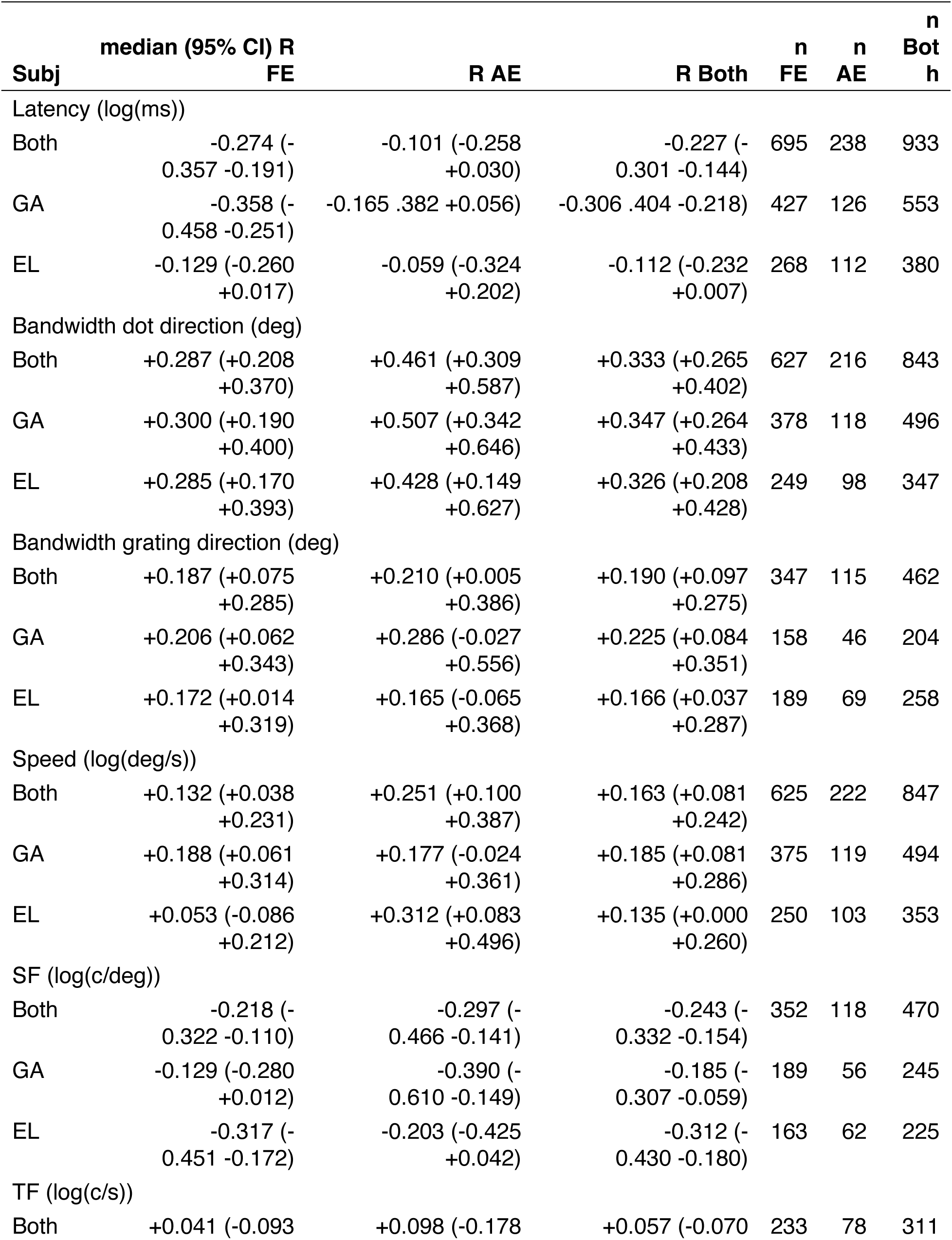
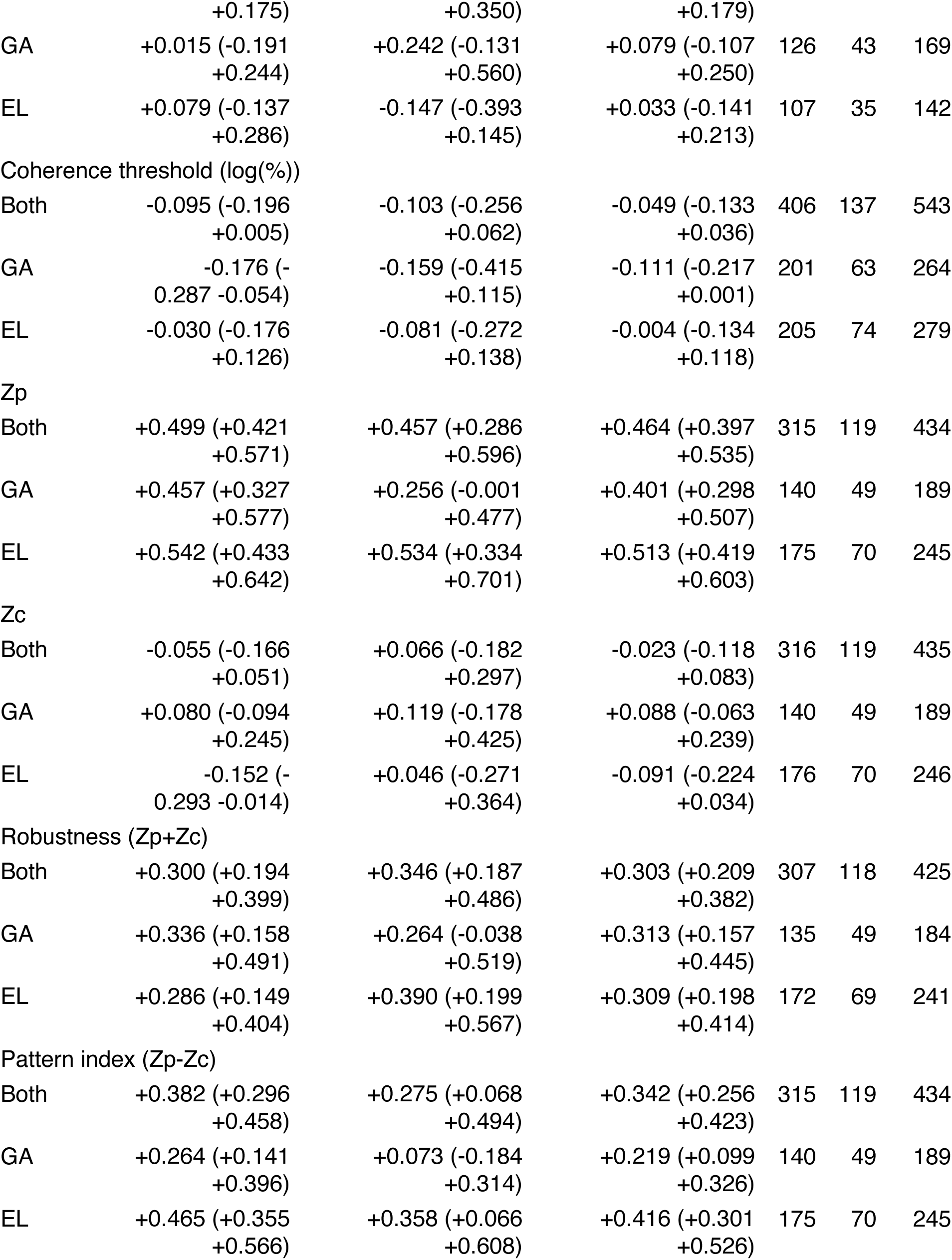
Correlation coefficients of tuning properties and response magnitude.

### Interocular suppression

Another possibility is that the scarcity of recorded AE cells could be a result of active suppression of the AE by the FE in the awake case. Evidence of active inhibition can be found in a small subset of recorded cells that we noticed because they showed spontaneous activity under blank or random noise conditions, but ceased firing when a stimulus was presented monocularly in the non-preferred eye, especially when the parameters matched the preference in the other eye. These *super monocular* cells were scored with *ODI < -1* or *ODI > +1* and are shown in the outer bins in figure 5B. Increased suppression from the FE onto the AE would not be surprising, but inhibition was sometimes also observed when we stimulated through the FE. Although this is a small subset of cells, they have an interesting implication. Somehow the super monocular cell is inhibited without visual stimulation in the preferred eye. These results could be explained by an active mechanism that acts to eliminate disruptive signals from the AE. One might reason that the amblyopic visual system, faced with conflicting signals from the two eyes, simply chooses to suppress one and work with the other.

### Spatiotemporal characteristics of MT neurons

Behaviorally, our animals, like amblyopic humans (Meier et al., 2016), showed lower than normal sensitivity for slow speeds. We found that this behavioral loss is reflected in neural sensitivities as well. Our results are corroborated by findings in anesthetized amblyopic animals reported in El-Shamayleh et al. (2010). However, there was little effect of amblyopia on temporal dynamics in general. We found no significant difference in temporal frequency tuning, and the difference in speed tuning that we observed between amblyopic and fellow eye neuron populations seems to be secondary to significant tuning differences in spatial frequency. Unlike the dearth of AE neurons we identified, these spatial frequency deficits could be inherited from upstream visual areas such as V1. We have previously observed reduced spatial resolution in V1 of amblyopic macaques (Movshon et al. (1987); Kiorpes et al. (1998)).

### Response latency and reliability

Although behaviorally, amblyopic observers tend to have slower reaction times and saccadic latencies when viewing through the amblyopic eye (Mackensen, 1958; Niechwiej-Szwedo et al., 2010, 2012; McKee et al., 2016), we found that amblyopic neurons in fact have *shorter* response latency. An onset latency difference for amblyopic neurons compared to those from visually typical monkeys has been reported before, with amblyopic latency being shorter than normal (Wang et al., 2017). However, in that case recordings were from ventral stream area V2 in amblyopic and normal control monkeys, and no direct comparison was made between fellow and amblyopic eye neurons. Interestingly, El-Shamayleh et al. (2010) reported shorter integration times for amblyopic eye neurons suggesting that overall response dynamics may be altered in amblyopia in a way that makes the AE faster.

The amblyopic visual system is sometimes characterized as “noisy” (e.g., Levi, Klein and Chen, 2008; Wang et al., 2017), therefore it is important to consider the reliability of neural responses driven through each eye. We found that AE neuron responses showed greater variability than FE responses, although these effects were modest, and smaller than the differences in Fano factor noted by El-Shamayleh et al. (2010). In our analysis, we found a fluctuation of response gain over time that is likely to be responsible for increased variability and decreased reliability of amblyopic eye responses. Further, Wang et al. (2017) reported substantially greater overall variability, trial-to-trial variability, and variable spontaneous activity in amblyopic V2, suggesting that indeed amblyopic extrastriate cortex may be “noisier” in general than visually typical cortex.

## Conclusion

In conclusion, this study provides an extensive characterization of neural organization in amblyopic area MT in macaques. Although previous studies were conducted exclusively under anesthesia, we found good correspondence between this work and our previous study and indeed corroborated our findings by subsequent recording of our awake behaving subjects under anesthesia. The dramatic loss of responsiveness of the amblyopic eye neural population, coupled with specific losses in coherence sensitivity and a shift to faster speed ranges provide strong correlates to behaviorally measured deficits in global motion sensitivity.

## Acknowledgements

We thank Michael Gorman, the members of the Visual Neuroscience Laboratory, and the staff of the NYU Office of Veterinary Resources for their assistance in rearing, behaviorally testing, and caring for the animals. Najib Majaj and Andrew Zaharia gave valuable assistance during surgery and awake recordings, and Paul Levy helped with the acute recording experiments. This research was supported by grants from the National Eye Institute to L.K. (R01 EY05864) and to L.K. and J.A.M. (R01 EY24914).

