## Supplementary figures for "Responses to visual motion of neurons in the extrastriate visual cortex of macaque monkeys with experimental amblyopia"

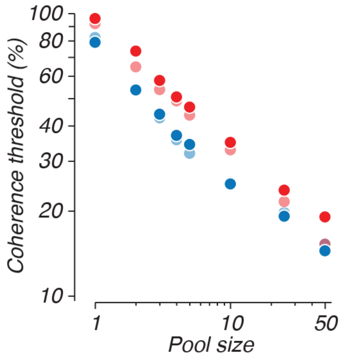


**Supporting figure 1. Effect of pool size on the simulated coherence thresholds of populations of neurons.** In red are pools created with AE cells, in blue FE cells. Light-colored pools are cells from subject GA. EL is dark-colored. While changing the pool size the fractional differences between pools are maintained, e.g. when doubling the pool size, the difference in threshold are divided by two. The choice of pool size therefore does not affect the qualitative differences observed between FE and AE, which show that for all pool sizes the FE coherence threshold is lower.

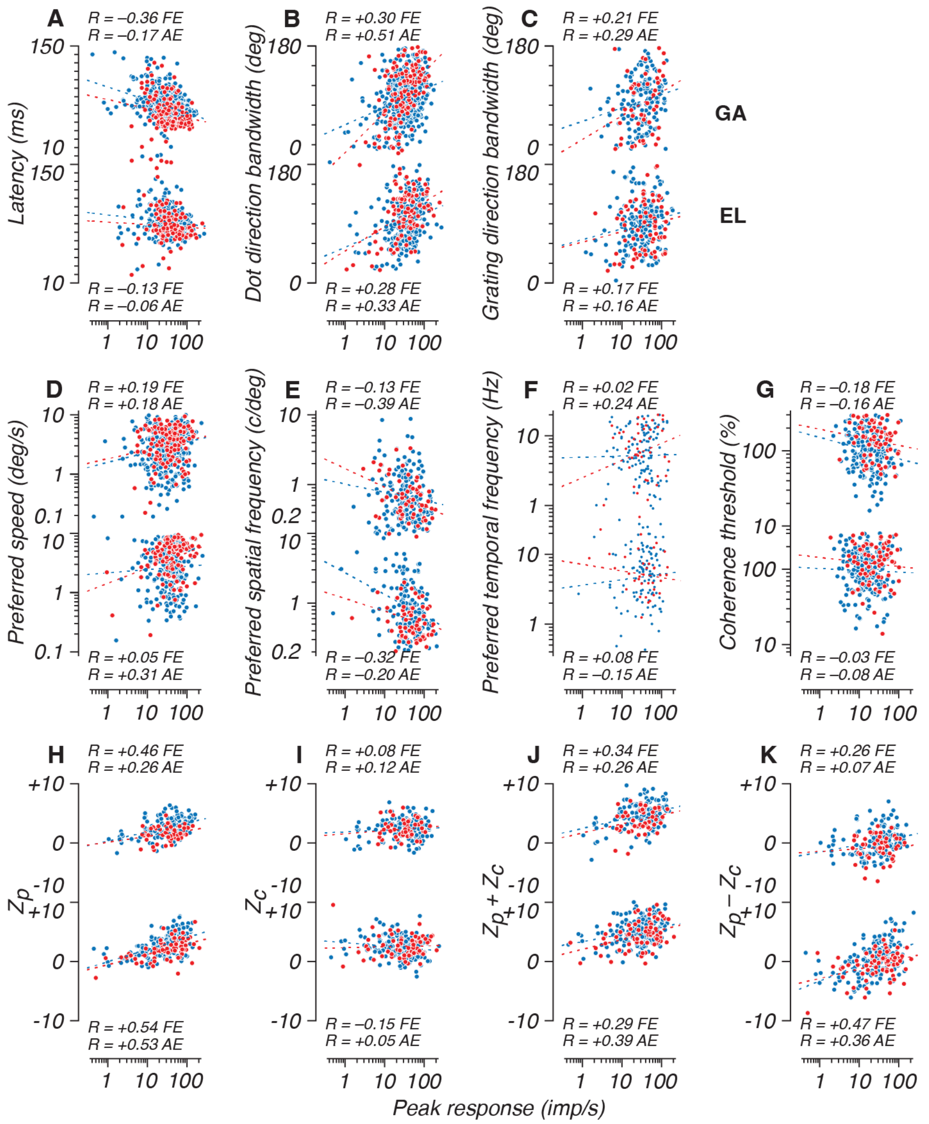


**Supporting figure 2. Regression of peak response and tuning properties**. Dashed lines are best fitting lines. Pearson correlation coefficients (R) of tuning properties against response magnitude are shown for FE and AE and for subject GA (top) and EL (bottom). See also Table 2 and Figure 12.
